# Senescent fibroblasts in the tumor stroma rewire lung cancer metabolism and plasticity

**DOI:** 10.1101/2024.07.29.605645

**Authors:** Jin Young Lee, Nabora Reyes, Sang-Ho Woo, Sakshi Goel, Fia Stratton, Chaoyuan Kuang, Aaron S. Mansfield, Lindsay M. LaFave, Tien Peng

**Author notes:** Address correspondence to: Tien Peng, M.D., University of California, San Francisco 513 Parnassus Ave., HSE Building, Room 1312 San Francisco, CA 94143.

## Abstract

Senescence has been demonstrated to either inhibit or promote tumorigenesis. Resolving this paradox requires spatial mapping and functional characterization of senescent cells in the native tumor niche. Here, we identified senescent *p16^Ink4a^*+ cancer-associated fibroblasts with a secretory phenotype that promotes fatty acid uptake and utilization by aggressive lung adenocarcinoma driven by Kras and p53 mutations. Furthermore, rewiring of lung cancer metabolism by *p16^Ink4a^*+ cancer- associated fibroblasts also altered tumor cell identity to a highly plastic/dedifferentiated state associated with progression in murine and human LUAD. Our *ex vivo* senolytic screening platform identified XL888, a HSP90 inhibitor, that cleared *p16^Ink4a^*+ cancer- associated fibroblasts *in vivo*. XL888 administration after establishment of advanced lung adenocarcinoma significantly reduced tumor burden concurrent with the loss of plastic tumor cells. Our study identified a druggable component of the tumor stroma that fulfills the metabolic requirement of tumor cells to acquire a more aggressive phenotype.

## Introduction

Senescence presents a paradox for how multicellular organisms maintain proliferative homeostasis over their lifespan, as this age-related cellular process has been shown to either attenuate or promote tumorigenesis^1^. On one hand, the induction of cell cycle arrest driven by known tumor suppressors would support senescence’s role as a crucial checkpoint against malignant transformation. However, Judith Campisi demonstrated in 2001 that senescent human lung fibroblasts promoted tumor growth when co-cultured together *in vitro*, and this phenomenon is driven by secreted factors from senescent fibroblasts^2^. While the ability of senescent fibroblasts to drive tumorigenesis has been replicated over time *in vitro*, it is not clear if this recapitulates a functional mechanism of tumor progression *in vivo*. The challenge is to spatially resolve the identity of senescent cells in the tumor niche and characterize their direct interaction with tumors *in vivo*.

*KRAS* and *TP53* are two of the most prevalent mutations found in lung adenocarcinoma (LUAD), the occurrence of which also correlates with advanced staging and shorter survival. Genetic mouse model combining KrasG12D mutation (K) with p53 deletion (P) produces a more aggressive LUAD that is relatively resistant to standard chemotherapy when compared to either mutations alone^3^. The induction of KP also generates more intratumoral heterogeneity when compared with K alone, as single cell studies have highlighted unique tumor subsets that arise from advanced KP-LUAD^4,5^. These emergent tumor subsets, arising from mature alveolar type 2 cells (AT2), are marked by gene regulatory programs characterized by a dedifferentiated or “plastic” cell state that recapitulates the lineage of foregut endoderm from which AT2 originates. Also of note is that the dedifferentiated tumors share markers with recently identified transitional cell states arising from AT2 to alveolar type 1 (AT1) differentiation during injury repair^6–8^, which is modified by stromal factors in the AT2 niche^6^. These data suggest that reorganization of the tumor stroma could play a vital role in the emergence of LUAD subsets that drive disease progression.

Leveraging our ultrasensitive reporter of *p16^Ink4a^* we previously demonstrated that senescent fibroblasts in healthy tissues promote epithelial stem cell regeneration after injury^9^. In this study, we set out to determine whether cancer co-opts the regenerative properties of senescent cells in a transformed stem cell niche. By spatial mapping of *p16^Ink4a^*+ cells in the tumor stroma and characterizing their interactions with LUAD, we aim to resolve the senescence paradox by deconstructing the tumor-intrinsic and tumor- extrinsic role of senescence in cancer.

## Results

### Emergence of *p16^Ink4a^*+ cancer-associated fibroblasts in the tumor stroma in aggressive LUAD

To characterize *p16^Ink4a^*+ cells during malignant transformation *in vivo*, we crossed our <u>INK</u>4A H2<u>B</u>-GFP <u>R</u>eporter-In-<u>T</u>and<u>e</u>m (INKBRITE) mouse^9^ with an autochthonous model of aggressive LUAD (Kras^G12D/+^;Trp53^fl/fl^;Rosa26^tdTomato/+^)^3^. This combined reporter/LUAD model (hereafter referred to as KPTI) enabled simultaneous tracing of tumor (tdTomato+) and *p16^Ink4a^*+ cells (nuclear GFP+) after adenoviral-mediated Cre recombination (**Fig. 1A**). We observed extensive infiltration of GFP+ cells within the tumor stroma around 8-10 weeks following the induction of LUAD in the KPTI mice *via* intratracheal adenoviral Cre delivery (**Fig. 1B**). Immunohistochemistry (IHC) analysis of the tumor demonstrated that many of the GFP+ cells within the tumor are ACTA2+, which is a cancer-associated fibroblast (CAF) marker associated with myofibroblastic differentiation (myCAF), whereas GFP+ cell on the tumor margin co-localize with the inflammatory CAF(iCAF)/adventitial fibroblast marker, PI16(peptidase inhibitor16) (**Fig. 1C**)^10,11^. We performed single cell transcriptome analysis of sorted GFP+ (*p16^Ink4a^*+) and GFP- (*p16^Ink4a−^*) fibroblasts from KPTI lungs, then merged this data with our reference dataset of fibroblast populations from the normal lung. We utilized previously published gene signature of CAF populations to annotate our CAF populations^10,11^, which demonstrated that the majority of *p16^Ink4a^*+ fibroblasts in KPTI lungs are myCAFs, which are absent in the normal lung (**Fig. 1D, Supplemental Fig. 1A-C, Supplemental Table 1,2**). In contrast, the iCAF population mostly clustered with adventitial fibroblasts in the normal lung (hereafter referred to as iCAF/adventitial) (**Fig. 1D, Supplemental Fig. 1A-C**). Flow cytometry of myCAF-specific surface marker, ITGA1, confirmed that the majority of myCAFs were *p16^Ink4a^*+ (**Supplemental Fig. 1D,E**), and transcript analysis of *p16^Ink4a^*+ fibroblasts isolated from KPTI lungs demonstrated significant enrichment for myCAF markers (**Supplemental Fig. 1F**). Isolated *p16^Ink4a^*+ CAFs were highly enriched for characteristics associated with both senescent cells and myofibroblasts^12,13^, including F-actin aggregation, cell size enlargement, polynucleation, DNA damage (ψ-H2AX foci), proliferative arrest, and β-galactosidase activity (**Fig. 1E, Supplemental Fig. 1G-I**).

**Figure 1.**
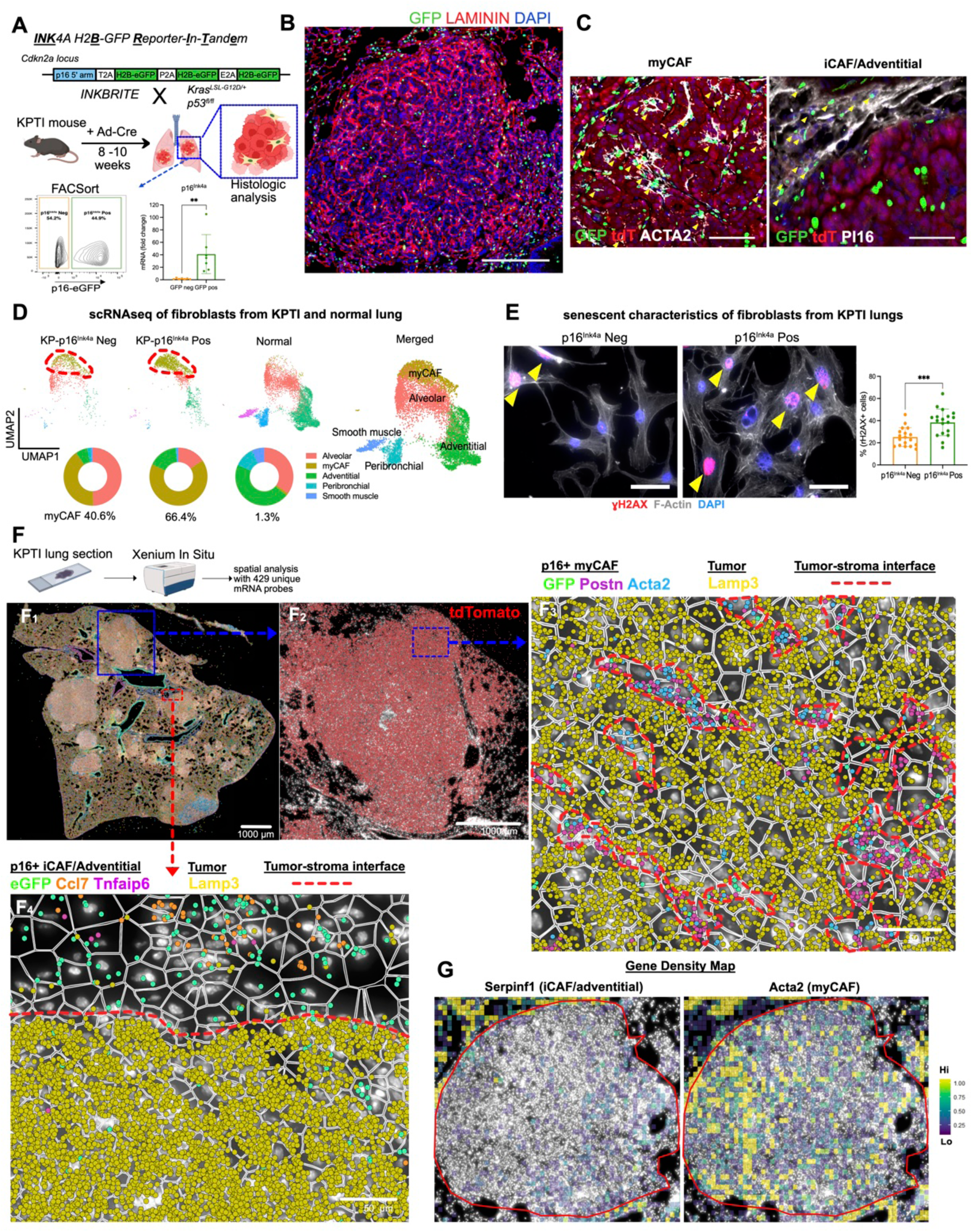
Contribution of senescent *p16^Ink4a^*+ fibroblasts to CAFs in mouse LUAD. (A) Schematic of experimental design to investigate *p16^Ink4a^*+ fibroblasts in mouse LUAD and demonstration of capacity to sort for *p16^Ink4a^*+ fibroblasts from LUAD tissues. (B) Representative immunofluorescence images showing nuclear GFP+ cells (green) within the stroma (Laminin+, red) in the lungs of KPTI mice at 8-10 weeks post-tumor induction. Scale bars, 200 μm. (C) Representative image of immunostaining of GFP, ACTA2, PI16, and tdTomato in the lungs of KPTI mice at 10 weeks post-tumor induction. Scale bars, 50 μm. (D) Top: UMAP plot of scRNA-seq data from fibroblasts isolated from KPTI and normal mouse lungs. Bottom: Proportion of fibroblast subtype relative to the total fibroblast population within each condition. (E) Left: Representative image of immunostaining for ɣH2AX and F-Actin in *p16^Ink4a−^* and *p16^Ink4a^*+ fibroblasts isolated from KPTI mouse lungs. Right: Quantitative analysis of ɣH2AX+ cells (n=18 per group). Scale bars, 50 μm. (F) Spatial profiling of mouse LUAD section using Xenium In Situ to elucidate the distribution of *p16^Ink4a^*+ fibroblasts expressing CAF markers within LUAD tissue. Each colored dots represents transcript detection overlaid on segmented cell borders. Tumor/stroma defined by presence/absence of tumor and stromal-specific transcripts. (G) Gene expression density map of iCAF/adventitial and myCAF markers relative to the tumor margins. Unpaired *t*-test was used in (E) to test statistical significance. Data are represented as mean ± SD.; \*\*\**P* < 0.001

To spatially resolve the gene signature associated with CAFs^10,11^ and recently identified LUAD subsets^4,5^, we conducted spatial analysis of 429 unique mRNA probes on section from a KPTI lung (12 weeks from induction) using the Xenium platform (**Fig. 1F1, Supplemental Table 3**). Probes for *tdtomato* and *GFP* enabled visualization of tumor and *p16^Ink4a^*+ cells respectively (**Fig. 1F2-4**). Overlaying spatial transcript coordinates (colored circles) with cellular segmentation, we could identify tumor stroma free of tumor transcripts (*Lamp3*) that were occupied by transcripts of *p16^Ink4a^*+ myCAFs (*GFP, postn, acta2*) (**Fig. 1F3**). In contrast, transcripts for *p16^Ink4a^*+ iCAFs/adventitial fibroblasts (*GFP, ccl7, tnfaip6*) were most localized on tumor margin (**Fig. 1F4**). This is confirmed by gene transcript density mapping demonstrating infiltration of myCAF markers within the tumor and iCAF/adventitial fibroblast markers on the tumor margins (**Fig. 1G**).

### *p16^Ink4a^*+ CAFs form a spatially segregated niche with an aggressive LUAD subset

We integrated single cell spatial data generated by Xenium into the Seurat workflow for clustering using uniform manifold approximation and projection (UMAP), followed by cell annotation using gene signature enrichment (UCell)^14^. We were able to identify clusters of previously identified CAFs^10,11^ along with LUAD subsets that ranged from cells with mature alveolar type 2 (AT2) markers to a dedifferentiated/plastic cell state called “high-plasticity cell state” (HPCS) identified in murine lungs with KRAS mutation^5^ (**Fig. 2A, Supplemental Fig. 2A, Supplemental Table 4**). The tumor clusters generated from the spatial data is comparable to the single cell analysis of sorted tdTomato+ cells from KPTI lungs (**Supplemental Fig. 2B-D, Supplemental Table 5**). Spatial analysis demonstrated that HPCS cells occupy distinct regions within the tumor, and are characterized by the induction of genes associated with alveolar transitional states emerging with injury (e.g. *Cldn4, Krt7, S100a14*)^6,7^ concurrent with the loss of canonical AT2 gene expression (*e.g. Lamp3, Sftpc, Hc*) (**Fig. 2B**, **Supplemental Fig. 2E**).

**Figure 2.**
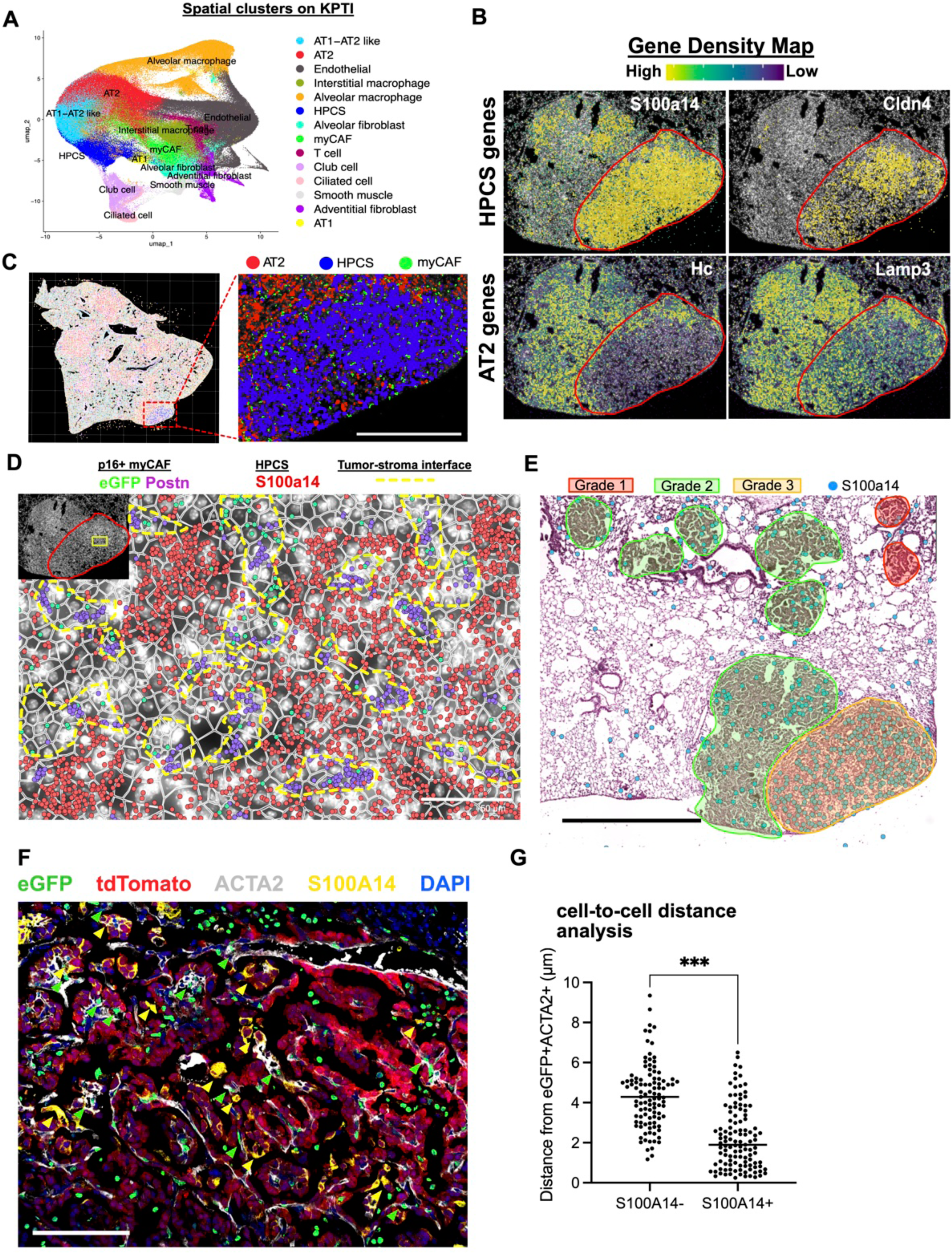
*p16^Ink4a^+* CAFs promote LUAD progression by supporting adjacent HPCS cells. (A) UMAP derived from spatially analyzed transcripts from KPTI mouse lung section. (B) Gene Density of HPCS and AT2 markers within the tumor. (C) ImageDimPlot of KPTI mouse lung with cell positions annotated by cluster labels, with localization of AT2, HPCS, and myCAF clusters in the region of interest. Scale bars, 500 μm. (D) Transcript mapping with cell segmentation highlighting the proximity between *p16^Ink4a^*+ myCAFs and S100a14+ HPCS cells. Scale bars, 50μm. (E) S100a14 transcript localization aligned with H&E of KPTI lungs with tumor histologic grading for aggressive features. Scale bars, 1000 μm. (F) Immunofluorescence identification of GFP+ACTA2+ fibroblasts (indicated by green arrowheads) and S100A14+tdTomato+ HPCS cells (indicated by yellow arrowheads) in KPTI mouse LUAD. Scale bars, 100 μm. (G) Quantification of the distance between GFP+ACTA2+ fibroblasts and S100A14+ or S100A14- tumor cells, with individual measurements represented as data points (n=103 for S100A14-, n=115 for S100A14+). Unpaired *t*-test was used in (G) to test statistical significance. Data are represented as mean ± SD.; \**P* < 0.05, \*\*\**P* < 0.001, \*\*\*\**P* < 0.0001

Mapping the distinct clusters generated by Seurat back onto the tumor section, we observed spatial segregation of HPCS cells within the tumor that are infiltrated by *p16^Ink4a^*+ myCAFs (**Fig. 2C,D**), and these areas of HPCS correlated with higher histologic tumor grades (**Fig. 2E**). Cell-to-cell distance analysis of tumor sections demonstrated that *p16^Ink4a^*+ myCAFs (GFP+/ACTA2+, green arrows) are located closer to HPCS (S100A14+/tdTomato+, yellow arrows) than non-HPCS (S100A14-/tdTomato-) cells within the tumor (**Fig. 2F,G**). These results demonstrate that *p16^Ink4a^*+ myCAFs are preferentially localized adjacent to an aggressive LUAD subset, suggesting a functional interaction between the tumor stroma and specific tumor subsets that is driving tumor heterogeneity and progression.

### *p16^Ink4a^*+ CAFs promote the emergence of aggressive LUAD subset

To explore the functional role of p16^Ink4a+^ CAFs in LUAD progression, we established a 3D organoid co-culture system on an air-liquid interface using freshly sorted fibroblasts (GFP+ vs. GFP-) and tumor cells (tdTomato+) from KPTI lungs (**Fig. 3A**). Coculture of *p16^Ink4a^*+ CAFs with tumor significantly enhanced organoid growth (**Fig. 3B**). Single cell analysis of LUAD arising in KPTI yielded surface markers for HPCS, including LY6A, that enabled flow sorting (**Supplemental Fig. 2E**). Sorted LY6A+ tumor displayed significantly enhanced growth (**Supplemental Fig. 3A**). Analysis of the tumor-CAF organoids demonstrated that p16^Ink4a^+ CAFs significant increased LY6A+ tumors, which is confirmed on flow analysis of the tumor organoids (**Fig. 3C,D**). Transcript analysis of sorted tumor cells (tdTomato+) confirmed the upregulation of HPCS markers concurrent with downregulation of AT2 markers in tumor organoids with *p16^Ink4a^*+ CAFs (**Supplemental Fig. 3B,C**). We performed single cell RNAseq analysis of the tumor organoids with gene signature enrichment analysis for cellular annotation, which confirmed the increase in HPCS subsets when cocultured with *p16^Ink4a^*+ CAFs *in vitro*, and demonstrated recapitulation of the LUAD heterogeneity subsets seen *in vivo* (**Fig. 3E,F, Supplemental Fig. 3D, Supplemental Table 6**). Finally, adoptive transfer of *p16^Ink4a^*+ CAFs along with tdTomato+ LUAD into wild-type recipient lungs resulted in a significant increase in tumor burden as quantified by whole-lung microCT when compared to tumors transferred with *p16^Ink4a−^* CAFs (**Fig. 3G,H**). Mirroring the findings from tumor organoids, *p16^Ink4a^*+ CAFs increased the population of HPCS cells expressing LY6A and S100A14 in the engrafted tumor (**Fig. 3I,J**). These experiments demonstrated that *p16^Ink4a^*+ CAFs are sufficient to drive the emergence of HPCS from LUAD, likely through secreted factors from the senescent CAFs.

**Figure 3.**
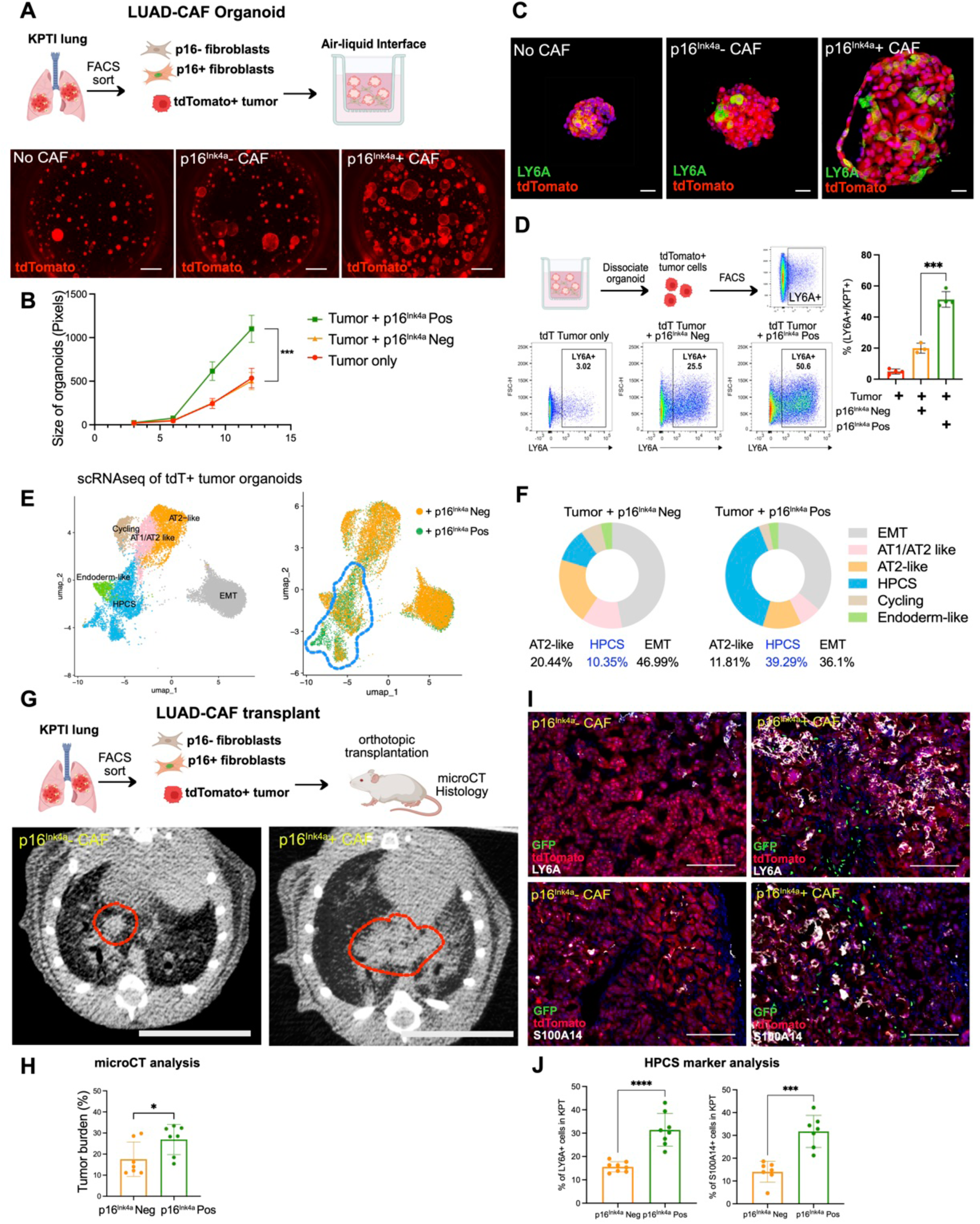
p16^Ink4a+^ fibroblasts support LUAD growth by increasing HPCS cells *in vitro* and *in vivo*. (A) Top: Schematic of the co-culture setup of FACS-sorted fibroblasts and tdTomato+ LUAD cells under air-liquid interface conditions. Bottom: Images of 3D tumor organoids. (B) Quantitative analysis of organoid sizes established in (A). (C) Representative images of LY6A immunofluorescence in organoids formed as described in (A). Scale bars, 100 μm. (D) Flow cytometry analysis of LY6A+ cell populations in 3D tumor organoids. (E) Single-cell RNA sequencing of tdTomato+ cells from organoids established in (A). Left: Unsupervised clustering of scRNA-seq data, annotated based on Marjanovic et al. Right: UMAP plot showing distinct cellular population that emerges in the tumor organoids co-cultured with p16^Ink4a+^ fibroblasts. (F) Pie graph depicting the proportion of cells contributing to identified clusters. (G) Top: Outline of the *in vivo* transplantation of tdTomato+ LUAD cells with either p16^Ink4a+^ or p16^Ink4a–^ fibroblasts into NSG mice. Bottom: microCT images of mouse lungs 4 weeks post-transplantation, Scale bars, 1 cm. (H) Tumor burden quantification in transplanted mice, expressed as tumor volume relative to whole lung volume at 4 weeks post-transplantation. (I) Representative image of S100A14 and LY6A immunofluorescence in the lungs of recipient mice, 4 weeks post-transplantation. Scale bars, 100 μm. (J) Quantitative analysis of HPCS cell prevalence within tdTomato+ tumor lesions. Unpaired *t*-test was used in (B), (D), (H), (J) to test statistical significance. Data are represented as mean ± SD.; \**P* < 0.05, \*\*\**P* < 0.001, \*\*\*\**P* < 0.0001

### *p16^Ink4a^*+ CAFs secrete APOE to promote highly plastic LUAD phenotype

We applied NicheNet^15^, an algorithm that predicts ligand-receptor interactions in single cell data, to predict interactions of ligands which are highly expressed in *p16^Ink4a^*+ CAFs with cognate receptors expressed in LUAD from our single cell analysis of KPTI lungs (**Fig. 4A**). Our analysis identified several ligands highly expressed in *p16^Ink4a^*+ CAFs and corresponding receptors on tumor cells. Among these ligands, APOE protein was markedly upregulated in *p16^Ink4a^*+ myCAFs (**Fig. 4B**), and *Apoe* transcript was significantly enriched in *p16^Ink4a^*+ CAFs (**Fig. 4C**). Spatial transcriptomic analysis confirmed the presence of *Apoe* transcripts in *p16^Ink4a^*+ myCAFs (**Fig. 4D**). Addition of recombinant APOE protein to *p16^Ink4a−^* CAFs cocultured with LUAD recapitulated the effects of *p16^Ink4a^*+ CAFs in tumor organoid model, with APOE enhancing organoid growth (**Fig. 4E,F**) while increasing the fraction of LY6A+ LUAD (**Fig. 4G-H**).

**Figure 4.**
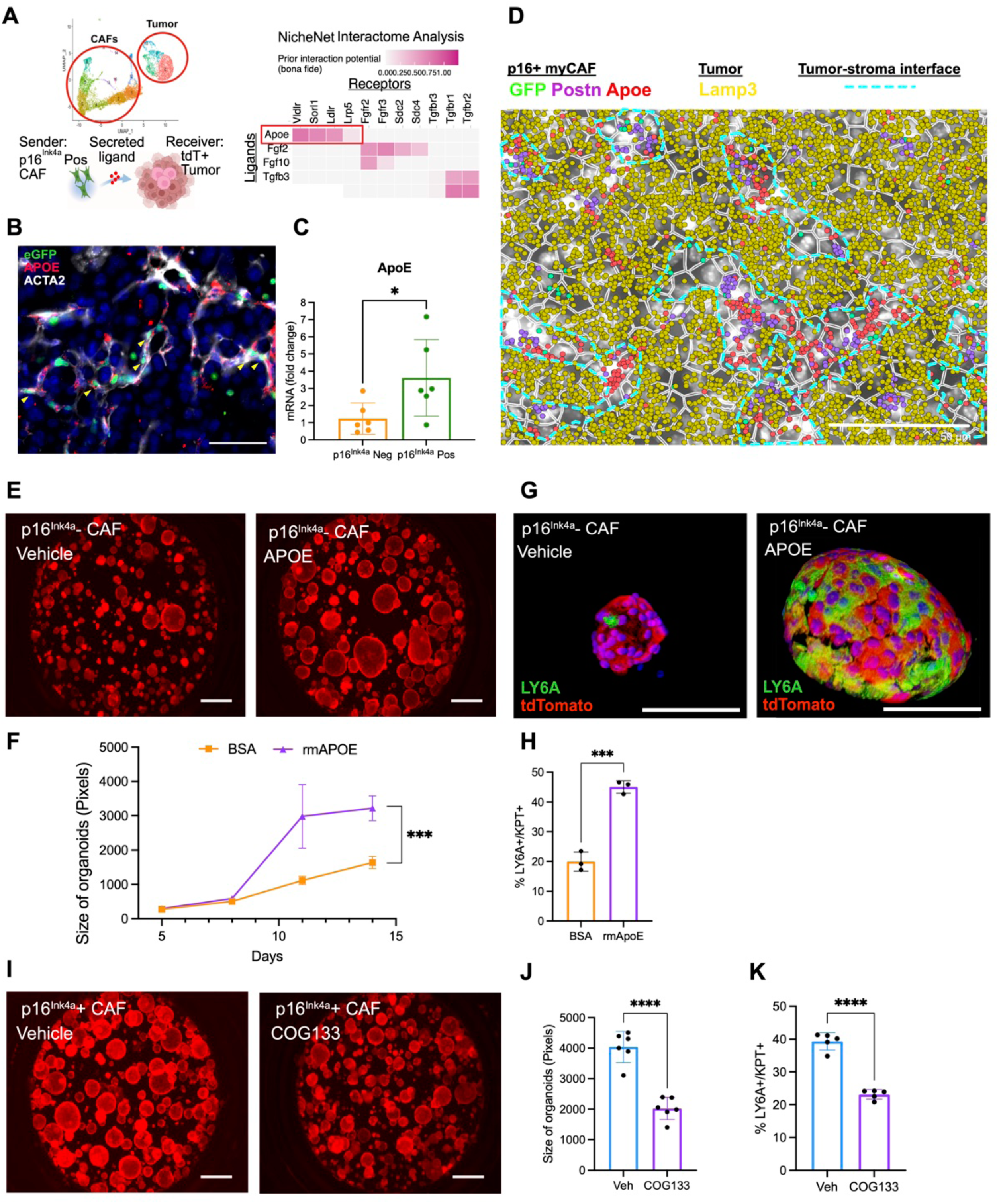
APOE derived from p16^Ink4a+^ fibroblasts promotes LUAD expansion by enriching the HPCS population. (A) NicheNet analysis of ligand-receptor interactions between p16^Ink4a+^ fibroblasts and tdTomato+ LUAD cells from KPTI mouse lungs. (B) Representative image of APOE immunofluorescence in KPTI lung tissue, indicating high expression of APOE in GFP+ACTA2+ fibroblasts. (C) qPCR evaluation of Apoe expression in fibroblasts sorted from KPTI mouse lungs. Each data point represents a separate biological replicate. (D) Xenium in situ visualization of Postn, Apoe, and GFP expression patterns within tumor areas in KPTI mouse lung. (E) Images of 3D tumor organoids treated with recombinant mouse APOE (rmAPOE). (F) Assessment of tumor organoid sizes with rmAPOE treatment. (G) Immunofluorescence detection of LY6A in tumor organoids derived from the experiment in (E). Scale bars, 100 μm. (H) Determination of LY6A+ cell percentages in organoids via flow cytometry. (I) Representative image of 3D tumor organoids treated with COG133, an ApoE mimetic peptide, to observe inhibitory effects on p16^Ink4a+^ fibroblasts function. (J) Measurement of sizes of tumor organoids established in (H). (K) Flow cytometry-based quantification of LY6A+ cells in tumor organoids treated with COG133. Unpaired *t*-test was used in (C), (F), (H), (J), and (K) to test statistical significance. Data are represented as mean ± SD.; \**P* < 0.05, \*\*\**P* < 0.001, \*\*\*\**P* < 0.0001

Conversely, blocking APOE binding to low density lipoprotein receptor (LDLR) through the APOE mimetic peptide, COG133, attenuated the effects of *p16^Ink4a^*+ CAFs in the tumor organoids by decreasing proliferation and the emergence of HPCS cells (**Fig. 4I- K**).

### APOE secreted by *p16^Ink4a^*+ CAFs promote lipid uptake and utilization by HPCS

APOE is a pleiotropic extracellular protein best known for its role in lipid transport^16^. We noticed that the KPTI tumor contained areas with vacuolated cytoplasm suggestive of lipid-laden cells. Oil red O and Bodipy staining confirmed lipid-laden tumor cells with HPCS markers (**Fig. 5A**), and spatial transcriptomic analysis demonstrated enrichment of *Apoe* and HPCS transcripts in the vacuolated regions of tumor (**Fig 5B, Supplemental Fig. 4A**). Aggressive cancer cells rely on fatty acids as a fuel source for biosynthesis and energy^17,18^. Bodipy stain of LUAD organoid treated with APOE or cocultured with *p16^Ink4a^*+ CAFs demonstrated increase in lipid droplet formation in LY6A+ tumor (**Fig. 5C, Supplemental Fig. 4B**). Addition of Bodipy-conjugated long chain fatty acid (LCFA) to the tumor organoids treated with APOE demonstrated that HPCS cells increased LCFA uptake (relative to non-HPCS cells) at baseline that is significantly upregulated in the presence of APOE (**Fig. 5D,E**). We knocked down *Apoe* expression in *p16^Ink4a^*+ CAFs (Apoe KD) using lentiviral-shAPOE (**Supplemental Fig. 4C**), which resulted in reduced LCFA uptake in the tumor organoids cocultured with Apoe KD *p16^Ink4a^*+ CAFs (**Fig. 5F**) and reduced HPCS in the tumor organoid (**Supplemental Fig. 4D).** LCFA profiling of the tumor organoids by liquid chromatography-mass spectrometry demonstrated an enrichment of LCFA 18-24 carbons in length in both tumor co-cultured with p16^Ink4a^+ CAFs and those treated with APOE (**Fig. 5G, Supplemental Fig. 4E**). Inhibition of the fatty acid β-oxidation (FAO) pathway (**Fig. 5H**) utilizing the CPT1 inhibitor, etomoxir, led to a significant reduction in proliferation along with markedly decreased HPCS in the tumor organoid cocultured with *p16^Ink4a^*+ CAFs (**Fig. 5I-K**). These results show that the senescent tumor stroma can rewire lipid metabolism of aggressive LUAD through APOE to increase tumor fitness.

**Figure 5.**
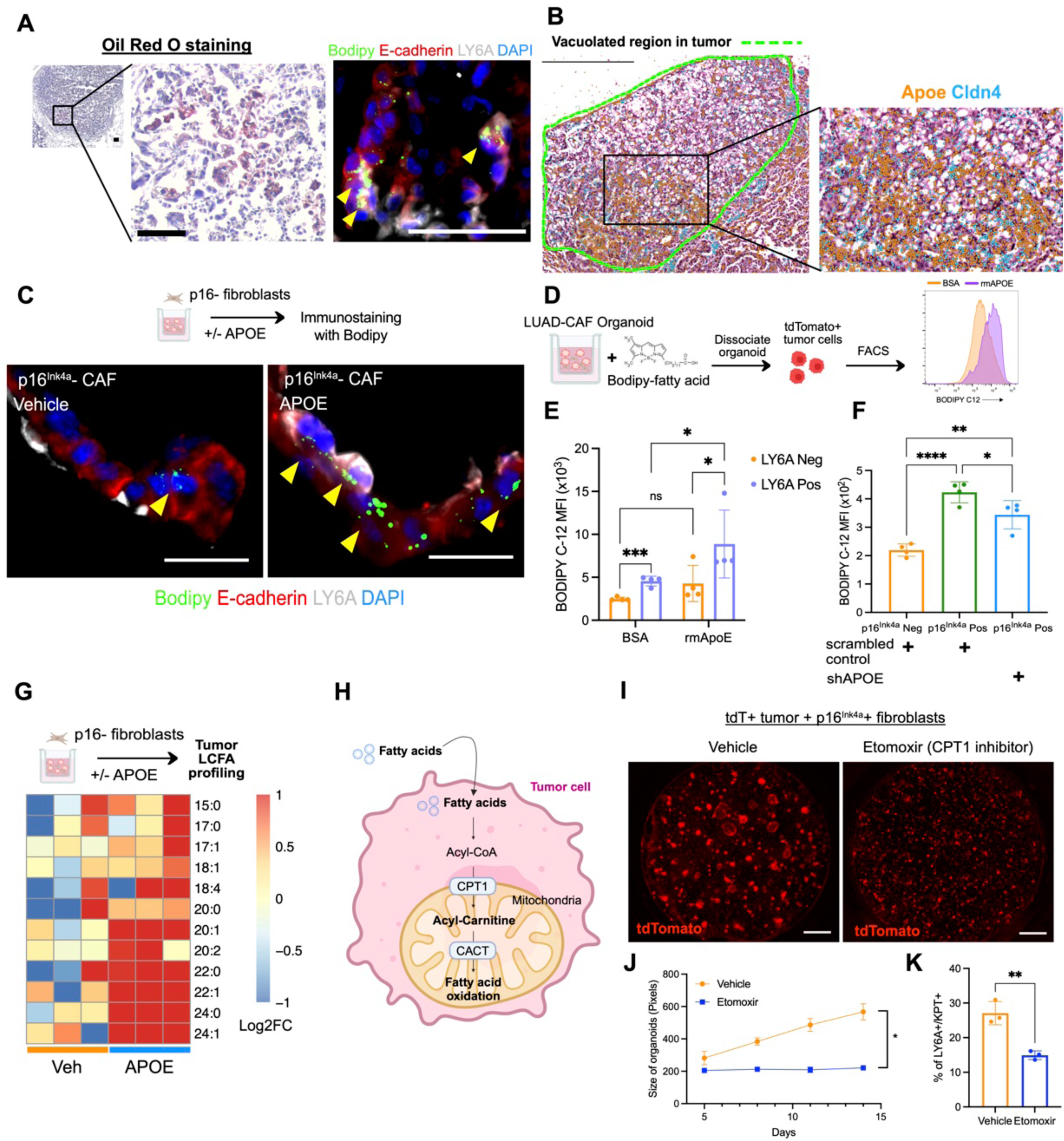
APOE from p16^Ink4a+^ fibroblasts modulates lipid metabolism in LUAD. (A) Left: Oil Red O staining of KPTI mouse lung tissue for lipid deposits. Scale bars, 100 μm. Right: LY6A immunofluorescence coupled with Bodipy 493/503 staining for neutral lipids in KPTI mouse lung. Scale bars, 50 μm. (B) Xenium in situ imaging aligned with H&E staining demonstrates localization of Apoe and Cldn4 transcripts within vacuolized regions in KPTI lung tissue. (C) Top: Schematic of 3D tumor organoid culture with treatment of rmAPOE protein. Bottom: Representative images of immunofluorescence of LY6A and Bodipy 493/503 staining for neutral lipids in 3D tumor organoids. Scale bars, 25 μm. (D) Diagram outlining the Bodipy fatty acid (Bodipy-C12) transfer assay in 3D tumor organoids. (E) Graph illustrating the increase in Bodipy-C12 Mean Fluorescence Intensity (MFI) in tdT+ tumor cells following rmAPOE treatment, suggesting enhanced fatty acid uptake. Right: MFI quantification of Bodipy-C12 in tdT+ tumor cells co-cultured with p16^Ink4a+^ fibroblasts (F) MFI quantification of Bodipy-C12 in tdT+ tumor cells co-cultured with p16^Ink4a+^ fibroblasts; downregulation of Apoe using shRNA indicates the role of APOE in supporting tumor cell fatty acid uptake. (G) Heatmap showing the log2 fold change in free fatty acid levels in tumor organoids cultured with p16^Ink4a–^ fibroblasts, comparing the effects of vehicle and rmAPOE treatment. (H) Schematic representing the metabolism of fatty acids, including their conversion into acyl-carnitines via carnitine palmitoyltransferase 1 (CPT1) for entry into fatty acid oxidation pathways. (I) Images of tumor organoids co-cultured with p16Ink4a+ fibroblasts and treated with either vehicle or etomoxir, a CPT1 inhibitor. (J) Organoid size quantification reveals that inhibition of fatty acid oxidation disrupts the tumor-supportive role of p16^Ink4a+^ fibroblasts. (K) Flow cytometry analysis of the LY6A+ cell proportion in tumor organoids subjected to fatty acid oxidation inhibition. One-way ANOVA was used in (E) and (F) and unpaired *t*-test was used in (J) and (K) to test statistical significance. Data are represented as mean ± SD.; \**P* < 0.05, \*\**P* < 0.01, \*\*\**P* < 0.001, \*\*\*\**P* < 0.0001

### Clearance of *p16^Ink4a^*+ CAFs suppresses tumor progression

Prior single cell analyses have demonstrated high similarity in transcriptomes of myCAFs arising in cancer and myofibroblast/fibrotic fibroblast enriched in TGFβ activation in fibrotic tissues^11,19^, which is consistent with our data when merging single cell datasets of *p16^Ink4a^*+ fibroblasts isolated from LUAD and lung fibrosis (bleomycin) models (**Supplemental Fig, 5A-C, Supplemental Table 7**). We recently reported a platform to screen for senolytic compounds targeting fibrotic *p16^Ink4a^*+ fibroblasts in lung fibrosis utilizing precision cut lung slice (PCLS) culture^20^. To screen for multiple senolytic compounds for efficacy against *p16^Ink4a^*+ CAFs within the preserved tumor microenvironment, we generated high-volume PCLS cultures from KPTI lungs (**Fig. 6A,B**) that enabled quantification of GFP+/GFP- fibroblast ratio by FACS after compound treatment *ex vivo* (**Fig. 6C**). Our lead senolytic compound in the lung fibrosis screen was XL888, a heat shock protein 90 inhibitor^20^. In KPTI-PCLS, XL888 significantly reduced GFP+ fibroblast fraction after 5 days of treatment as analyzed by FACS (**Fig. 6D**), which is confirmed on IHC analysis of the PCLS showing clearance of p16^Ink4a^+ myCAFs (GFP+/ACTA2+) within the intact tumor stroma (**Fig. 6E,F**). FACS analysis of KPTI-PCLS also demonstrated that BH3 mimetics^21^ (navitoclax and ABT- 737) have potential efficacy in clearing *p16^Ink4a^*+ CAFs (**Supplemental Fig. 5D**).

**Figure 6.**
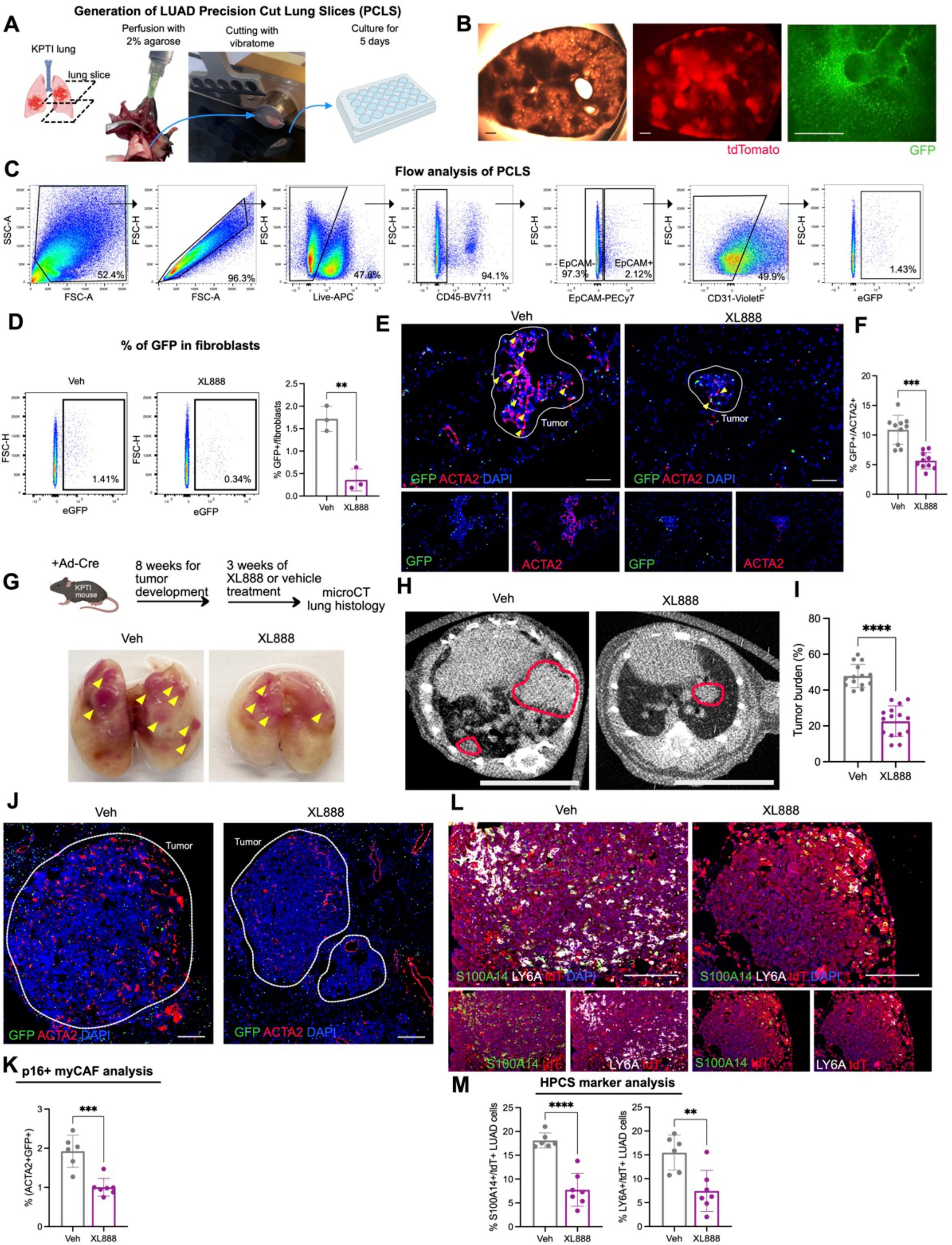
Senolytic compound XL888 clears p16Ink4a+ fibroblasts and reduces tumor burden in KPTI mice. (A) Generation of PCLS cultures from KPTI mouse lungs. (B) Brightfield and fluorescence images of PCLS demonstrating the preservation of tdTomato+ LUAD and GFP+ cells. Scale bars, 1000 □m. (C) Flow cytometry strategy for evaluating GFP+ cells within sorted (CD45-EpCAM- CD31-) fibroblasts post-XL888 treatment. (D) Flow cytometry analysis of GFP+ fibroblasts in PCLS treated with XL888 (n=3 slices for each group). (E) Representative image of GFP and ACTA2 immunofluorescence in PCLS treated with vehicle or XL888. Scale bars, 100 □m. (F) Quantitative analysis of GFP+ cells among ACTA2+ fibroblasts (n=10 for each group). (G) Experimental design to test tumor suppressive effect of XL888 in KPTI mouse (top), alongside representative macroscopic lung images from vehicle- and XL888-treated groups (bottom). (H) MicroCT images of lungs from KPTI mice following treatment. Scale bars, 1 cm. (I) Assessment of tumor burden in KPTI mice by microCT, expressed as the tumor volume to whole lung volume ratio (n=14-15 mice for each group). (J) Representative image of GFP and ACTA2 immunofluorescence of KPTI mouse lung tissue. Scale bars, 200 □m. (K) Quantification of p16Ink4a+ myCAFs in KPTI moues lung (n=6-7 mice for each group). (L) Representative image of S100A14, LY6A, and tdTomato immunofluorescence of KPTI lung tissue. Scale bars, 200 □m. (M) Quantification of the proportion of HPCS cells, identified by S100A14 (left) and LY6A (right), within the tdTomato+ LUAD cell population in KPTI mouse lungs (n=6-7 mice for each group). Unpaired t-test was used in (d), (f), (i), (k), and (m) to test statistical significance. Data are represented as mean ± SD.; **P < 0.01, ***P < 0.001, ****P < 0.0001

We then tested the effects of XL888 *in vivo* by treating KPTI mouse with XL888 after establishment of tumor 8 weeks out from induction (**Fig. 6G**). MicroCT quantification of tumor volume demonstrated that XL888 significantly reduced tumor burden after 3 weeks of treatment (**Fig. 6H,I**). IHC analysis confirmed the reduction in *p16^Ink4a^*+ myCAFs after XL888 treatment (**Fig. 6J,K**). Furthermore, XL888-treated KPTI lungs demonstrated decreased HPCS cells in the tumors (**Fig. 6L,M**). Flow cytometry analysis revealed no change in the percentage of *p16^Ink4a^*+ immune cells (GFP+/CD45+) in the XL888-treated KPTI lungs (**Supplemental Fig. 5E**). The PCLS senolytic screen demonstrated that the combination of dasatnib and quercetin (D&Q)^22^ did not have efficacy in clearing p16^Ink4a^+ myCAFs (**Supplemental Fig. 5D**). D&Q did not reduce *p16^Ink4a^*+ myCAFs nor tumor burden when administered *in vivo* to KPTI animals with established tumor (**Supplemental Fig. 5F-I**). These results suggested that the efficacy of senolytics is context/target dependent, and demonstrated that clearance of *p16^Ink4a^*+ myCAFs with XL888 attenuated tumor progression and prevented the emergence of plastic tumor subsets in an aggressive model of LUAD.

### *p16^INK4A^* promotes myCAF phenotype in human lung fibroblasts

Analysis of previously published cell atlas of CAFs identified in patients with non-small cell lung cancer (NSCLC)^23^ demonstrated the enrichment of *CDKN2A* (encodes both *p16^INK4A^*and *p14^ARF^* in alternative reading frames) in myCAFs that are also enriched for *CTHRC1* and *POSTN,* similar to the murine myCAFs (**Supplemental Fig. 6A**). To confirm the spatial localization of myCAFs in relation to tumor subsets in human LUAD, we applied our spatial analysis workflow utilizing a human probe set for lung epithelial and fibroblast subtypes (**Supplemental Table 8**) on a biopsy sample of LUAD with KRAS driver mutation (G12A). Transcript identification with cell segmentation generated unique clusters that were annotated using gene enrichment analysis and mapped onto the slide (**Fig. 7A,B, Supplemental Fig. 6B, Supplemental Table 9**). We were able to identify CDKN2A+/APOE+ myCAFs in the tumor stroma adjacent to recently described transitional/plastic cell (KRT8+/CLDN4+) found primarily in patients with KRAS-mutated LUAD^24^ (**Fig. 7B**). IHC analysis of LUAD samples confirms the presence of p16^INK4A^ protein in the myCAFs within the tumor stroma (**Supplemental Fig. 6C**). Applying a gene signature derived from single cell data of human *p16^INK4A^*+ myCAFs in NSCLC^23^ to the Cancer Genome Atlas (TCGA) dataset demonstrated a significant correlation between *p16^INK4A^*+ myCAF gene signature expression level in tumor samples and mortality in patients with LUAD (**Fig. 7C, Supplemental Table 10**).

**Figure 7.**
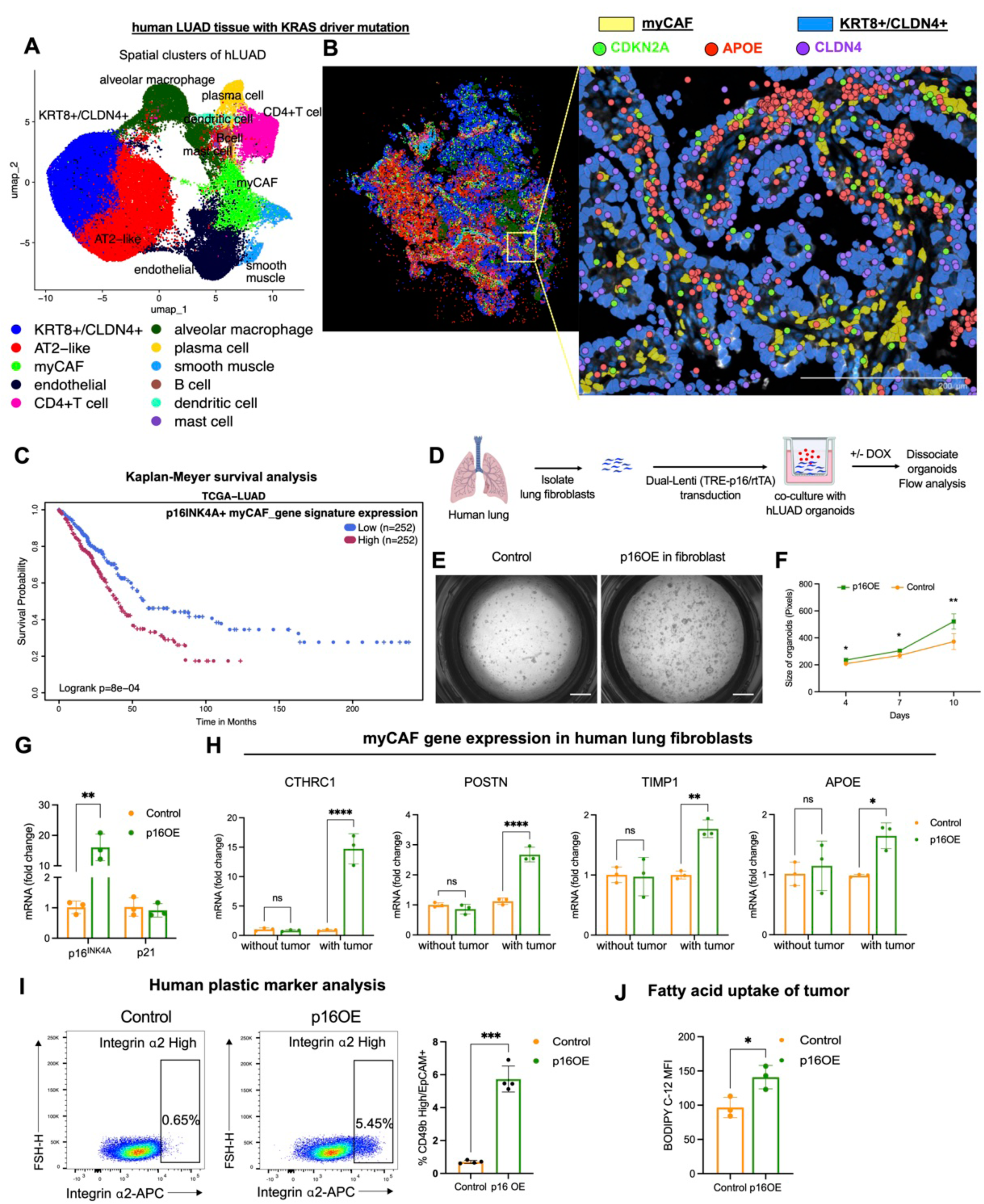
Contribution of *p16^INK4a^*+ fibroblasts to human LUAD progression. (A) UMAP derived from spatially analyzed transcripts from human LUAD section. (B) ImageDimPlot of human LUAD section with cell positions annotated by cluster labels (left), localization of KRT8+/CLDN4+ cells (blue), and myCAFs (yellow) with transcripts of CDKN2A, APOE, and CLDN4 in the region of interest (right). Scale bars, 200 μm. (C) Survival analysis of LUAD patients in the TCGA based on expression level of gene signature in the *p16^INK4A^*+ CAF cluster from a human CAF single cell data set (Grout *et al.*). (D) Experimental design to test the effects of *p16^INK4A^*-overexpression (p16OE) in human lung fibroblasts on human LUAD (hLUAD) organoids. (E) Images of hLUAD organoids cultured with either control or p16OE fibroblasts. Scale bars, 2000 μm. (F) Quantificaiton of hLUAD organoids in (e) (n=5 for each group). (G) qPCR analysis of *p16^INK4A^* and p21 in normal human lung fibroblasts after overexpression of *p16^INK4A^*(n=3 for each group). (H) qPCR analysis of control or p16OE lung fibroblasts with or without co-culture with tumor (n=3 for each group). (I) Left: Flow cytometry image of tumor organoids from (E). Right: Quantification of Integrin α2 high plastic cell population within tumor organoids (n=4 for each group). (J) Analysis of fatty acid transfer within tumor organoids established in (E) (n=3 for each group). Unpaired *t*-test was used in (F), (G), (I), (J) and and one-way ANOVA was used in (H) to test statistical significance. Data are represented as mean ± SD.; \**P* < 0.05, \*\**P* < 0.01, \*\*\**P* < 0.001, \*\*\*\**P* < 0.0001

To test the sufficiency of *p16^INK4A^* expression in the induction of the human myCAF phenotype, we designed a dual lentiviral system (Lenti-tTS/rTTA+Lenti-TRE- p16-2A-tdTomato) to overexpress (OE) *p16^INK4A^*in primary human fibroblasts isolated from control donors in a doxycycline (dox) dependent fashion. The transduced human lung fibroblasts are cocultured with human LUAD organoids generated from NSCLC patients with KRAS driver mutations like our 3D CAF-tumor organoid model for KPTI (**Fig. 7D**). Dox induction (p16OE) of the human CAF-tumor organoid significantly enhanced tumor growth *in vitro* (**Fig. 7E,F**). Transcript analysis of fibroblasts confirmed the induction of *p16^INK4A^*after dox treatment (**Fig. 7G**). Dox induction of *p16^INK4A^*in lung fibroblasts induced myCAF signature gene expression in fibroblasts cocultured with tumor, but not in the absence of tumor (**Fig. 7H**). This suggested that the tumor initiates the myCAF program in *p16^INK4A^*+ lung fibroblasts in a paracrine fashion, which then initiates a *p16^INK4A^*+ CAF-to-tumor signaling program to promote tumor progression.

Analysis of the tumor cells by flow cytometry demonstrated an increase in human plastic cell marker^5^, ITGA2, as well as an increase in Bodipy-LCFA uptake in the tumor cocultured with p16OE fibroblasts (**Fig. 7I,J**). Administration of human recombinant APOE variants to the tumor organoids demonstrated that hAPOE2 had the largest effect in promoting tumor organoid growth, which correlated with LCFA uptake in culture (**Supplemental Fig. 6D-F**).

## Discussion

One of the common links between Hayflicks’s original description of a cell cycle arrest termed “senescence”^25^ and Campisi’s seminal observation that senescent cells can drive malignant transformation^2^ is that both were studying human lung fibroblasts grown in culture. The significance of whether these phenotypes of lung fibroblasts *in vitro* correlate with their native function *in vivo* remains an open question. Our data demonstrate that senescent lung fibroblasts arising in the tumor stroma drive tumor evolution to a more dedifferentiated cellular state with altered metabolic requirement for high-energy substrate. These tumor-intrinsic features correlate with the known features of LUAD progression such as increased intratumoral heterogeneity^4,5,26^ and rewired fatty acid metabolism^17,18^, but we now present a model where tumor-extrinsic signals in the stroma can drive these cancer hallmarks.

The emergence of lineage heterogeneity in LUAD would also suggest divergent metabolic requirements that predicate their distinct function. Metabolic profiling of different lung cancer cell lines demonstrated remarkable diversity, even when cultured under standard conditions, that correlated with distinct oncogenic mutations as well as therapeutic response^27^. One of these divergent metabolic features is an increased dependence on fatty acid utilization to sustain energy production and biomass synthesis in certain tumor subtypes. Activation of KRAS has been demonstrated to increase both fatty acid uptake and synthesis in LUAD that drives tumor proliferation^17,28^. However, it was not clear whether the tumor microenvironment is playing a role in this divergent metabolic requirement for LUAD to utilize lipid as fuel, nor do we understand how lipid utilization differs amongst LUAD tumor subsets with varying capacity to drive progression. Our study demonstrated that senescent CAFs within the tumor stroma rewire the metabolic requirement of an aggressive LUAD subset, and this metabolic rewiring can drive tumor plasticity and sustain intratumoral heterogeneity as the LUAD progresses to more advanced stages.

We have previously demonstrated that senescent fibroblasts in the lung stem cell niche dynamically alter their secretory program to promote stem cell renewal during acute injury^9^. Conceptualizing cancer as a nonhealing wound, our data suggests that cancer coopts the adaptive properties of the senescent stroma in the lung to select for tumor subsets with enhanced capacity to propagate. In this view, senescent fibroblasts are neither good nor bad, but rather fulfills the specific requirement of the epithelial niche (normal or malignant) to drive stem cell renewal regardless of the costs.

Uncovering this requirement does reveal a therapeutic opportunity to target the tumor stroma using senolytics, but it should be noted that the senescent fibroblasts we identified driving stem cell renewal in acute injury and cancer respectively are different types of fibroblasts. The need for target specificity in selecting senolytics becomes more apparent as the heterogeneity of senescent cells *in vivo* becomes increasingly recognized.

## Acknowledgements

We thank Parnassus Flow Cytometry Core for assistance with cell sorting for bulk and single cell RNA analysis (P30DK063720). We thank Youngho Seo (Preclinical microCT core) for processing of micro-computed tomography images. GEO accession number for raw RNA sequencing data is listed in Methods. This work is supported by NIH grants R01HL160895 and R01HL155622 and CIRM DISC0-14460 to T.P., the Tobacco-Related Disease Research Program (TRDRP) postdoctoral award T33FT6395 and Basic Science Research Program through the National Research Foundation of Korea (NRF) funded by the Ministry of Education (2020R1A6A3A03038781) to J.L., and Nina Ireland Program Award for human lung collection.

## Author Contributions

J.L. and T.P conceived the experiments and wrote the manuscript. J.L., N.R., S.W., S.G., F.S. performed the experiments, collected samples, and analyzed data. C.K. and A.S.M. collected human materials. L.M.L. provided input on the experiments and manuscript.

## Methods

### Key resource table

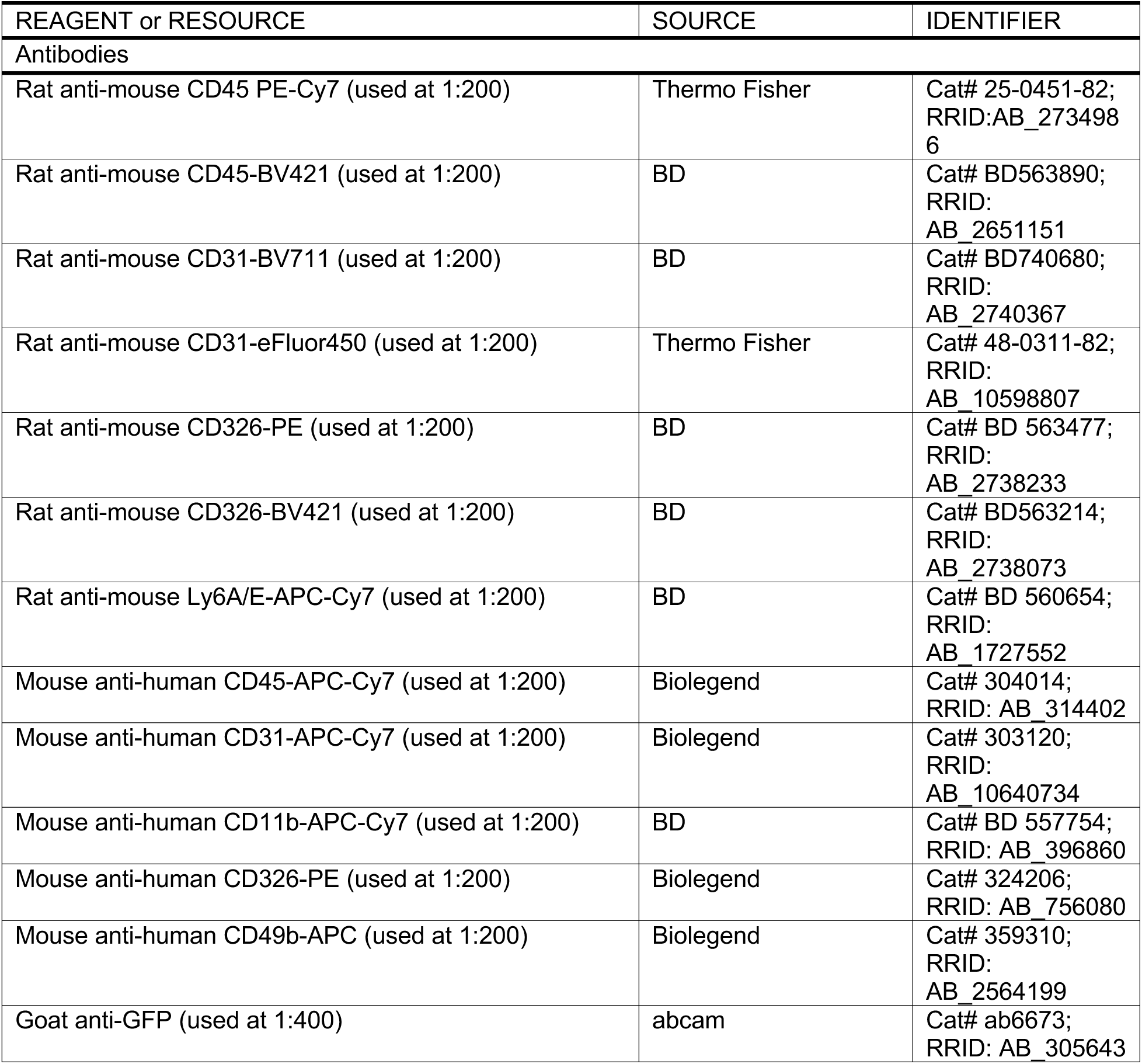
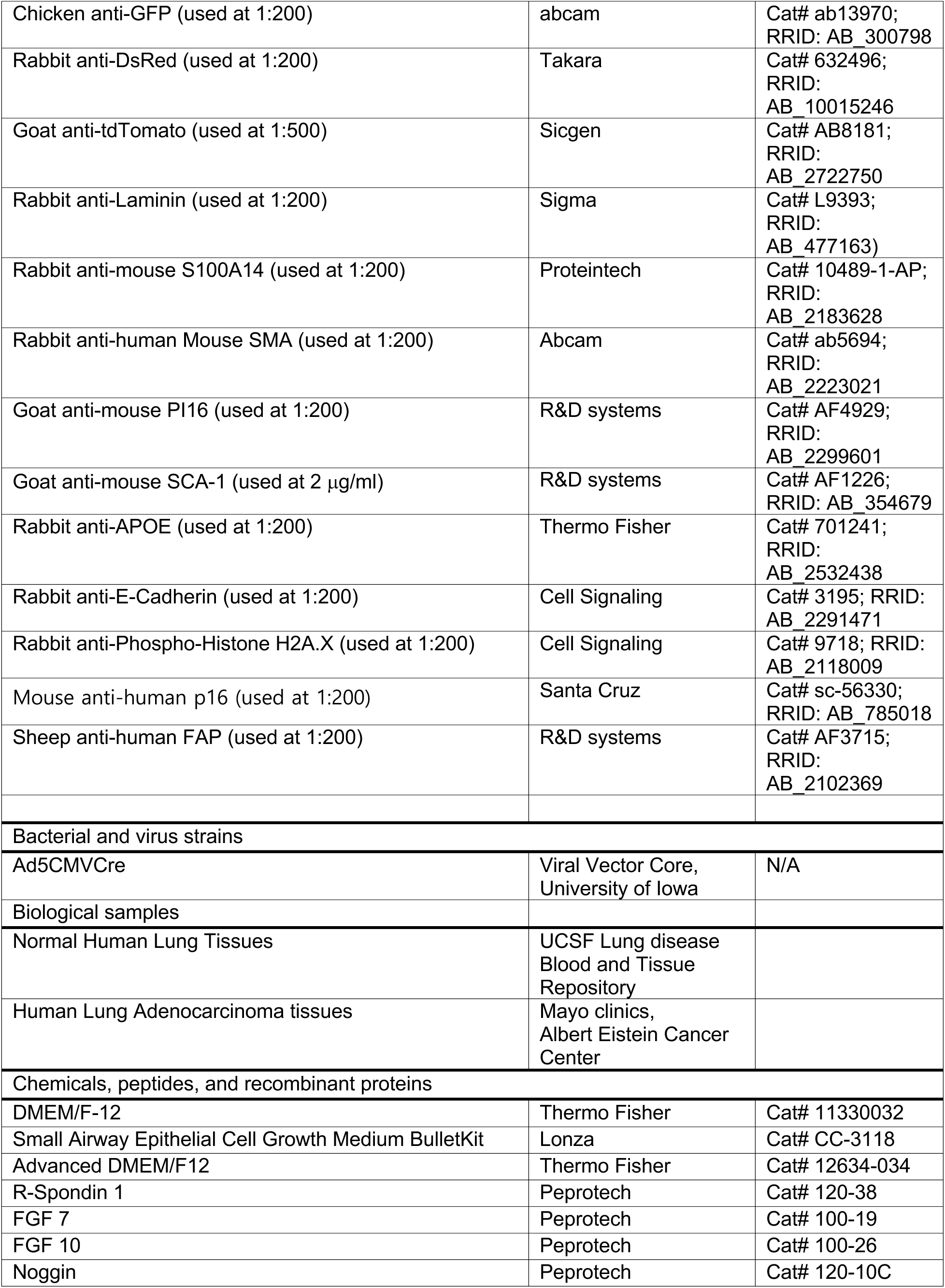
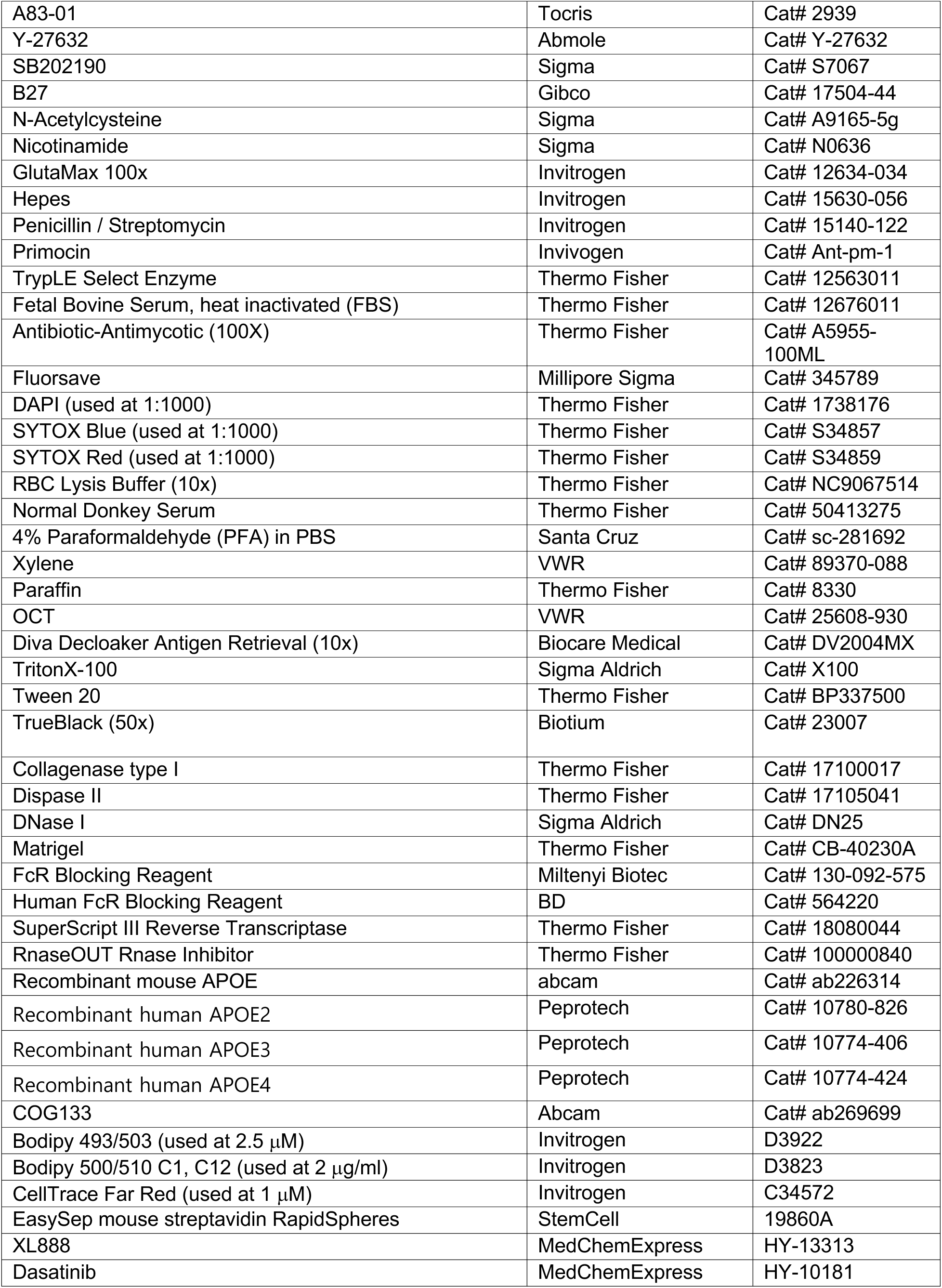
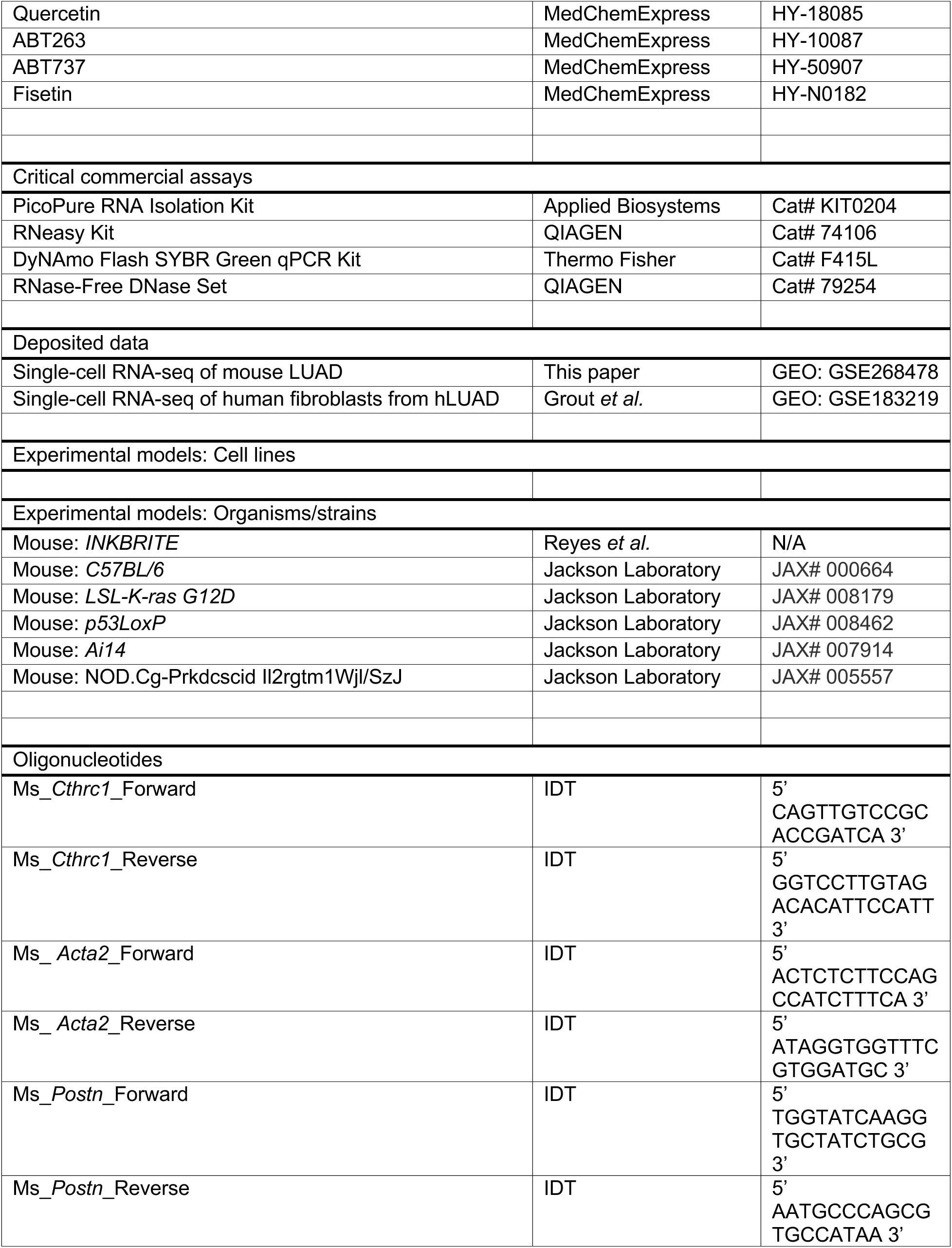
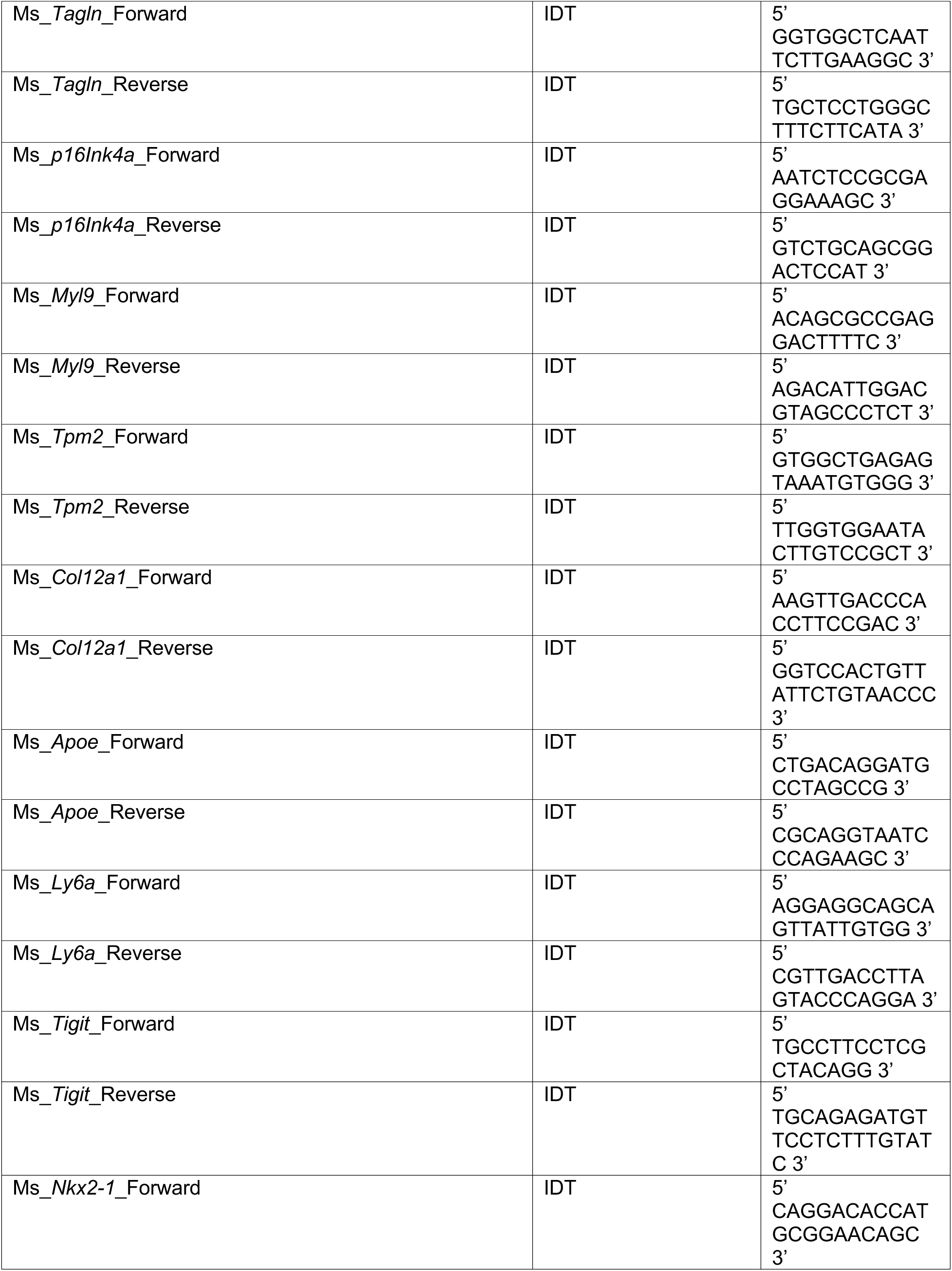
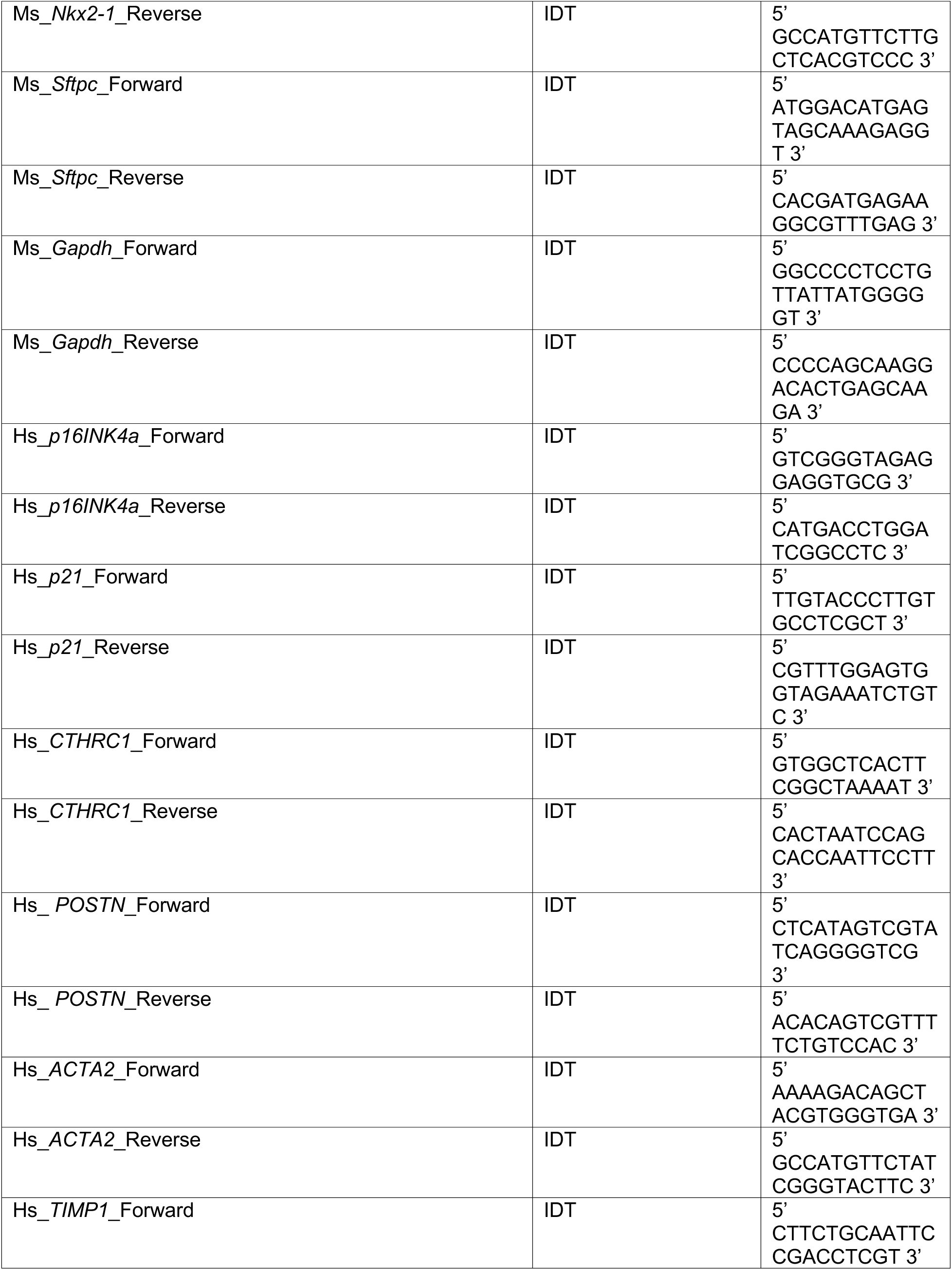
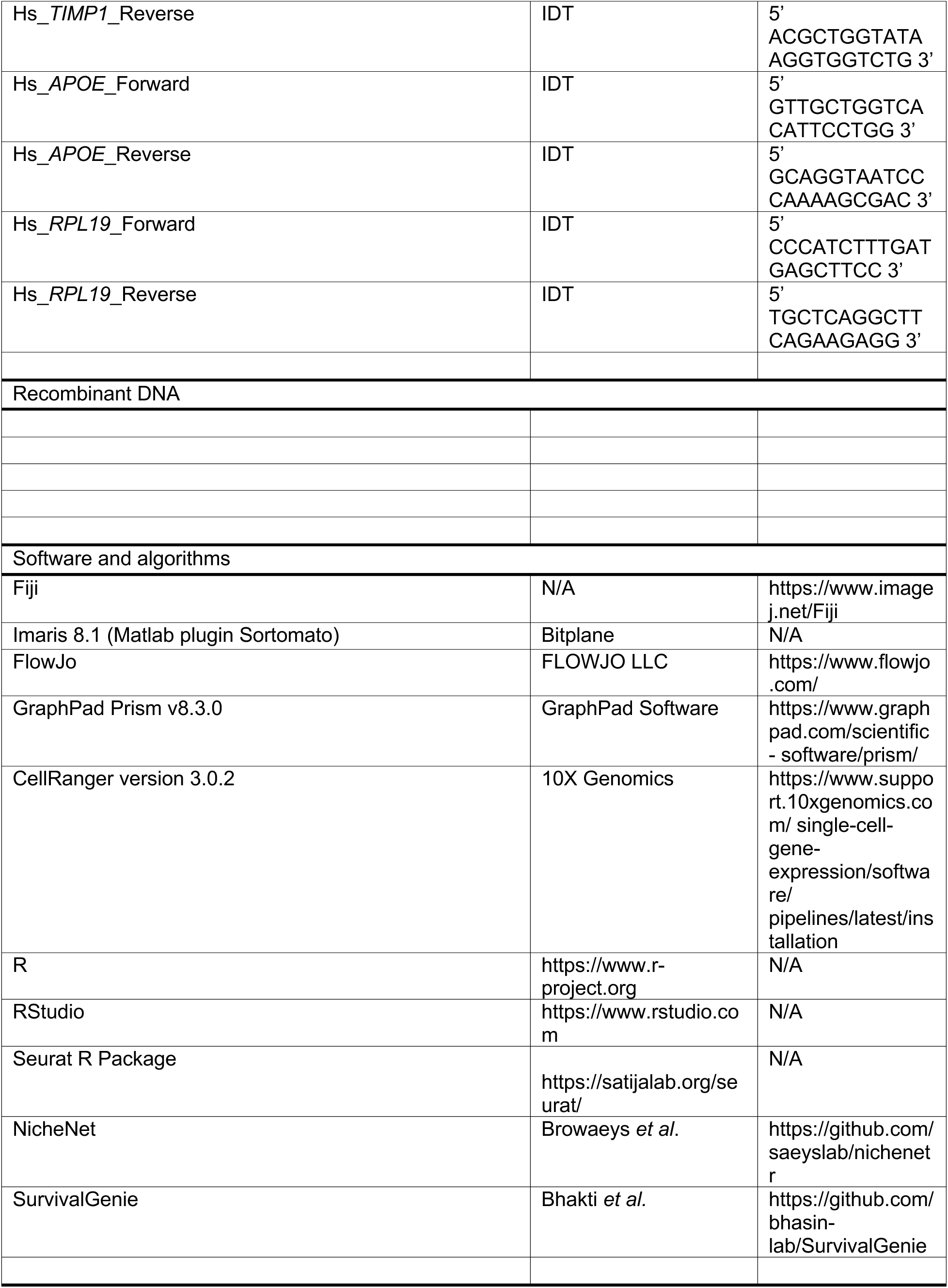
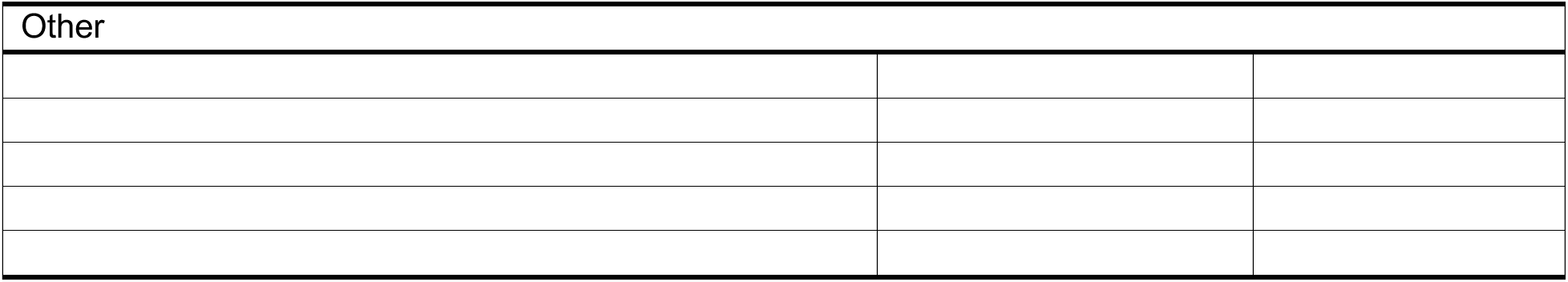

### Animal experiments

All animal studies described were approved by the IACUC at the University of California, San Francisco. All genetically engineered mice were maintained on a mixed or C57/BL6 background. Experiments were performed on male and female mice between 8-12 weeks old. Previously published *Kras^LSL-G12D/+^*^29^, *Trp53^flox/flox^* ^30^, *Rosa26^LSL-tdTomato/+^* ^31^, Apoe knock-out (JAX002052), and INKBRITE ^32^ mice were used in this study. In addition, we used NOD.Cg-Prkdc^scid^ Il2rg^tm1Wjl^/SzJ (NSG mice, The Jackson Laboratory, catalog #005557) in our transplantation studies. Tumors were induced in KP, KPT, or KPTI mice at 8 to 12 weeks of age with 2.5x10^7^ PFU of AdCMV-cre (University of Iowa) by intratracheal instillation as described previously ^33^. For XL888 treatment, KPTI mice were treated with XL888 as previously described ^34^. Briefly, 62.5 mg/kg of XL888 was delivered to mice via oral delivery 5 days a week for 3 weeks starting at 8 weeks after AdCMV-cre delivery. XL888 was dissolved in 10 mM hydrochloric acid with the concentration of 15.625 mg/ml. After vortexing, the dissolved XL888 was delivered to the mice using oral gavage daily. For dasatinib and quercetin (DQ) treatment, dasatinib (5 mg/kg) and quercetin (50 mg/kg) were administered to mice via oral gavage daily for 3 weeks, starting 6 weeks after adCMV-cre delivery. The compounds were dissolved in a solution consisting of 4% DMSO, 30% PEG300, 5% Tween80, 61% dH2O.

### Human Lung Tissue

Studies involving human tumor specimen were approved by the Mayo Clinic and Albert Einstein Institutional Review Board. All subjects provided written informed consent.

Human lung fibroblasts were isolated from the lungs of brain-dead donors that were rejected for lung transplantation. Clinical/demographic information of tissue donors are listed in Supplementary Table 11.

### FACS

Dissected mouse lung was tracheally perfused with a digestion cocktail of Collagenase Type I (225 U/ml, Thermo Fisher Scientific), Dispase (15 U/ml, Thermo Fisher Scientific) and Dnase (50 U/ml, Sigma-Aldrich) after perfusion with PBS. The lung was removed from the chest and incubated in a digestion cocktail for 45 minutes at 37 °C with continuous shaking. After digestion, remaining tissue chunks were finely minced with blades and washed with a FACS buffer (2% FBS and 1% Penicillin-Streptomycin in DMEM). The mixture was passed through a 70 μm cell strainer and resuspended in a red blood cell lysis buffer, then passed through 40 μm cell strainer. Cell suspensions were incubated with FcR blocker for 10 mins on the ice. After blocking, cell suspensions were incubated with the appropriate conjugated antibodies in a sorting buffer for 30 min at 4°C and washed with FACS buffer. Doublets and dead cells were excluded based on forward and side scatter and SYTOX Red (Invitrogen, S34859), respectively.

The following antibodies were used for staining: CD45-PE-Cy7 (Invitrogen, 50-112- 9643), CD45-BV421 (BD, 563890), CD31-BV711 (BD, 740680), CD31-BV421 (Invitrogen, 48-0311-82), EpCAM-PE (BD, 563477), EpCAM-BV421 (BD, 563214).

Immune (CD45-biotin, Biolgened, 103104), epithelial (CD326-biotin, Biolegend, 118204) and endothelial (CD31-Biotin, Biolegend, 102404) cells are removed with EasySep mouse streptavidin RapidSpheres (StemCell, 19860A), when applicable. FACS was performed on a BD FACS Aria using FACSDiva Software. CD45- CD31- EpCAM+ tdTomato+ cells were sorted for LUAD cells and CD45- CD31- EpCAM- cells were sorted for fibroblasts, the GFP- and GFP+ fibroblasts were further separated and were sorted into FACS buffer. Analysis was performed using FlowJo software.

For the human lung, a distal piece (∼10 cm^3^) was dissected from the whole lung and washed with HBSS X 4 times in 15 min. The piece of lung was further diced with razor blades and was added into the digestion cocktail of Collagenase Type I (225 U/ml, Thermo Fisher), Dispase (15 U/ml, Thermo Fisher) and Dnase (100 U/ml, Sigma). The mixture was incubated for 2 h at 37°C and vortexed intermittently. The mixture was then liquefied with a blender and passed through 4X4 gauze, a 100 mm and a 70 mm cell strainer. The mixture was resuspended in RBC lysis buffer, before passing through a 40 mm cell strainer. The cell suspensions were incubated with the antibodies in the FACS buffer for 30 min at 4°C and washed with the FACS buffer. The following antibodies were used: CD45-APC-Cy7 (BioLegend, 304014), CD31-APC-Cy7 (BioLegend, 303120), CD11b-APC-Cy7 (BD Biosciences, 557754), EpCAM-PE (BioLegend, 324206). DAPI (0.2 mg/ml) was used to exclude dead cells. Single cells were selected and CD45- CD11b- CD31- EpCAM- cells were sorted for fibroblasts. Cells were sorted into FACS buffer. FACS analysis was performed by FACSDiva (BD).

### Single-cell RNA sequencing and analysis

Single cell sequencing was performed on a 10X Chromium instrument (10X Genomics) at the Institute of Human Genetics (UCSF, San Francisco, CA). Briefly, live mouse lung cells were sorted and resuspended in 50 µl PBS with 0.04% BSA at 1,000 cells/µl and loaded onto a single lane into the Chromium Controller to produce gel bead-in emulsions (GEMs). GEMs underwent reverse transcription for RNA barcoding and cDNA amplification. The library was prepped with the Chromium Single Cell 3’ Reagent Version 3 kit. The samples were sequenced using the HiSeq2500 (Illumina) in Rapid Run Mode. We used the Seurat R package along with a gene-barcode matrix provided by CellRanger for downstream analysis. Following the standard workflow of Seurat, we generated Seurat objects after using ScaleData, RunPCA, RunUMAP. For human scRNA-seq data, we used processed scRNA-seq data from human LUAD fibroblasts from GSE183219. After generating subsets of lung fibroblasts, violin plots and density plots were generated.

### Xenium sample preparation

5 μm FFPE tissue sections from KPTI mouse lung tissues were placed onto a Xenium slide, followed by deparaffinization and permeabilization to make the mRNA detectable. The Xenium platform from 10X Genomics was used according to the manufacturer’s recommendations and as previously reported^35^. The Xenium output files were transferred for downstream analysis using the Xenium explorer.

### Post-Xenium histology

After running Xenium platform, H&E staining followed the manufacturer’s protocol (CG000160). Post-Xenium H&E images were registered to Xenium data using QuPath and Xenium explorer.

### Spatial cluster generation and mapping from Xenium

We employed the Seurat vignette (https://satijalab.org/seurat/reference/readxenium) to load and analyze the Xenium data with Seurat version 5. For normalization, we applied SCTransform method, followed by standard dimensionality reduction and clustering.

The clustering results were visualized in UMAP space. Subsequently, we annotated each cluster according to their gene expressions. The annotated clusters were imported into the Xenium explorer to map their spatial locations.

### Histologic grading of mouse tumors

Quantification of mouse lung tumor grade was performed by trained pathology technician who was blinded to sample ID. The quantification was based on parameters established by the previously described Aiforia platform that used automated deep neural network trained to classify NSCLC tumor grades (1-4) based on NSCLC_v25 algorithm^4^.

### Tumor transplantation experiment

8 weeks old recipient mice were injured by injecting 1-1.5U/kg bleomycin intratracheally 3 days before transplantation. Tumor cells and fibroblasts were obtained from LUAD- induced KPTI mice by FACS as detailed in section “FACS”. A mixture of 20,000 fibroblasts and 100,000 tumor cells or only tumor cells were resuspended in 50 μl PBS and introduced into the lungs of bleomycin-injured recipient mice intratracheally. After 4 weeks, lung tumor burden of recipient mice was evaluated using microCT imaging.

Subsequently, the mice were euthanized for lung histological analysis.

### 3D mouse tumor organoid cultures

Primary tumor organoid cultures were generated from tdTomato expressing tumor cells isolated from mice bearing 2-3 months old LUAD tumors. EpCAM+ tdTomato+ tumor cells were isolated by FACS and plated on Matrigel (Corning, CB-40230) as previously described ^32^. Briefly, tumor cells and GFP- or GFP+ fibroblasts were resuspended (4- 5x10^3^ tumor cells: 2.4-3x10^4^ fibroblasts/well) in 1:1 mixture of media and Matrigel. The media is comprised of small airway basal media (SABM) with selected components from SAGM bullet kit (Lonza) including Insulin, Transferrin, Bovine Pituitary Extract, Retinoic Acid, and human Epidermal Growth Factor. 0.1 µg/mL cholera toxin, 5% FBS, and 1% Penicillin-Streptomycin were also added. The mixture of cell suspension-matrigel-media was placed in a transwell of 24 well plate and allowed to solidify at 37°C. The growth media with 1 µM of ROCK inhibitor was added to the lower well of the well plate and refreshed with the media without ROCK inhibitor after 24 hr. Media was refreshed every 2-3 days.

### Human 3D tumor organoid cultures

Human 3D tumor organoid was established as previously described ^36^. Briefly, cryopreserved human LUAD specimen was thawed in 37°C and washed with PBS to remove freezing media. The tissue was placed in the plate with airway organoid media and minced with blade. The minced tissue was digested with digestion buffer (media + collagenase + Dispase + ROCK inhibitor) in 37°C shaker for 30 mins. The digested tissue was filtered with 100 µm filter and isolated cells were resuspended in Matrigel. 15-18 µl droplets of cell-Matrigel mixture was plated in 6-well plate. The cell-matrix mixture was allowed to polymerize for 20-30 mins and media was added to the well. The media was refreshed every 2-3 days.

### Histology and immunohistochemistry

For paraffin embedded mouse lungs, mouse right ventricles were perfused with 1 ml PBS and the lungs were inflated with 4% PFA, and then fixed in 4% PFA overnight at 4°C. After fixation, the lungs were washed by cold PBS X 4 times in 2 hrs at 4°C and dehydrated in a series of increasing ethanol concentration washes (30%, 50%, 70%, 95% and 100%). The dehydrated lungs were incubated with Xylene for 1 hr at RT and with paraffin at 65°C for 90 min X 2 times, and then embedded in paraffin and sectioned. Mouse PCLS samples were fixed in 4% PFA for 30 mins. After PBS washes, slices were embedded in OCT after 30% sucrose incubations. 5-8 μm thick cryosections were used for immunohistochemistry. Following antibodies were used: GFP (1:400, Abcam, ab6673), GFP (1:200, Abcam, ab13970), DsRed (1:200, Takara, 632496), tdTomato (Sicgen, 1:200, AB8181-200), RFP (Rockland, Rockland, 600-901-379), Laminin (Sigma, 1:200, L9393), S100A14 (Proteintech, 1:200, 10489-1-AP), Alpha smooth muscle actin (1:200, Abcam, ab5694), PI16 (R&D systems, AF4929), LY6A (R&D systems, 2 μg/ml, AF1226), APOE (1:200, Invitrogen, 701241), E-Cadherin (1:200, Cell signaling, 3195S), Phospho-Histone H2A.X (1:200, Cell signaling, 9718S) Human lung specimens were fixed and processed as the mouse lungs. Antibodies used for human lung slide staining were ACTA2 (1:200, Abcam, ab5694), p16INK4a (1:200, Santa Cruz, sc-56330), APOE (1:200, Invitrogen, 701241), FAP (1:200, R&D systems, AF3715). Images were captured using Zeiss Imager M1 or Leica Stellaris 5.

For organoids, the Matrigel containing organoids was fixed with 4% paraformaldehyde for overnight at 4°C. After multiple washes with PBS, the Matrigel was embedded in OCT for cryoblock.

### Neutral lipid droplet staining

Cryosections were used for the lipid staining with Bodipy 493/503 (Invitrogen, D3922). A 0.1% saponin-PBS solution was used for all the incubation buffers, including those for blocking and antibody staining. After secondary antibody staining, the sections were treated with 2.5 μM of Bodipy 493/503 for 30 mins at room temperature. Following PBS washes and DAPI staining, the slides were mounted and imaged.

### Organoid analysis

To obtain single cell suspensions from organoids, Matrigel containing organoids was digested with Dispase (15 U/ml, Thermo Fisher Scientific) and Dnase (50 U/ml, Sigma-Aldrich) for 30 mins at 37°C. Following the removal of Matrigel with Dispase, the samples were washed with PBS and treated with Triple (Gibco, 12604013) for 10 mins at 37°C to get single cells. The acquired single cells were then processed for FACS to get EpCAM+ LUAD cells or EpCAM- fibroblasts as described above.

### Cell Culture

Freshly isolated fibroblasts from KPTI lungs (GFP- or GFP+) or human lung fibroblasts were cultured in DMEM/F-12 (Thermo Fisher, 11330032) with 10% FBS and 1% Pen/Strep. The medium was changed every 2 days and lung fibroblasts were maintained for no more than 3 passages.

### CellTrace Far Red labeling (CTFR)

To compare proliferative capacity of fibroblasts, we utilized CellTrace Far Red cell labeling reagent (Invitrogen, C34572). Isolated fibroblasts were cultured for 3 days and then stained with CellTrace Far Red reagent. The fibroblasts were detached and stained with 1 μM of CellTrace for 20 minutes at 37°C (1 million cells per ml) following the manufacturer’s protocol. Post staining, the cells were washed with media and cultured for and additional 3 to 4 days. Serum-starved cells post-CTFR staining were used to separate CTFR high and low cells based on CTFR intensity levels. CTFR high cells, representing non-proliferating cells, were identified within the high intensity range of 95 to 97% encompassing serum-starve cells, while cells with lower intensity were considered as CTFR low cells.

### Quantitative RT-PCR (qPCR)

Total RNA was obtained from cells using PicoPure RNA Isolation Kit (Applied Biosystems, KIT0204) or RNeasy mini kit (QIAGEN, 74106), following the manufacturers’ protocols. cDNA was synthesized from total RNA using the SuperScript Strand Synthesis System (Thermo Fisher, 18080044). Quantitative RT-PCR (qRT-PCR) was performed using the SYBR Green system (Thermo Fisher, F415L). Relative gene expression levels after qRT-PCR were defined using the ΔΔCt method and normalizing to the housekeeping genes. The qRT-PCR primers used for mouse are as follows: Cthrc1-F: CAGTTGTCCGCACCGATCA; Cthrc1-R: GGTCCTTGTAGACACATTCCATT; Acta2-F: ACTCTCTTCCAGCCATCTTTCA; Acta2-R: ATAGGTGGTTTCGTGGATGC; Postn-F: TGGTATCAAGGTGCTATCTGCG; Postn-R: AATGCCCAGCGTGCCATAA; Tagln-F: GGTGGCTCAATTCTTGAAGGC; Tagln-R: TGCTCCTGGGCTTTCTTCATA; p16INK4a-F: AATCTCCGCGAGGAAAGC; p16INK4a-R: GTCTGCAGCGGACTCCAT; Myl9-F: ACAGCGCCGAGGACTTTTC; Myl9-R: AGACATTGGACGTAGCCCTCT; Tpm2-F: GTGGCTGAGAGTAAATGTGGG; Tpm2-R: TTGGTGGAATACTTGTCCGCT Col12a1-F: AAGTTGACCCACCTTCCGAC; Col12a1-R: GGTCCACTGTTATTCTGTAACCC; Apoe-F: CTGACAGGATGCCTAGCCG; Apoe-R: CGCAGGTAATCCCAGAAGC; Ly6a-F: AGGAGGCAGCAGTTATTGTGG; Ly6a-R: CGTTGACCTTAGTACCCAGGA; Tigit-F: TGCCTTCCTCGCTACAGG; Tigit-R: TGCAGAGATGTTCCTCTTTGTATC; Nkx2-1-F: CAGGACACCATGCGGAACAGC; Nkx2-1-R: GCCATGTTCTTGCTCACGTCCC; Sftpc-F: ATGGACATGAGTAGCAAAGAGGT; Sftpc-R: CACGATGAGAAGGCGTTTGAG; Gapdh-F: GGCCCCTCCTGTTATTATGGGGGT; Gapdh-R: CCCCAGCAAGGACACTGAGCAAGA The primers used for human are as follows: p16INK4a-F: GTCGGGTAGAGGAGGTGCG; p16INK4a-R: CATGACCTGGATCGGCCTC; p21-F: TTGTACCCTTGTGCCTCGCT; p21-R: CGTTTGGAGTGGTAGAAATCTGTC; CTHRC1-F: GTGGCTCACTTCGGCTAAAAT; CTHRC1-R: CACTAATCCAGCACCAATTCCTT; POSTN-F: CTCATAGTCGTATCAGGGGTCG; POSTN-R: ACACAGTCGTTTTCTGTCCAC; ACTA2-F: AAAAGACAGCTACGTGGGTGA; TIMP1-F: CTTCTGCAATTCCGACCTCGT; TIMP1- R: ACGCTGGTATAAGGTGGTCTG; APOE-F: GTTGCTGGTCACATTCCTGG; APOE-R: GCAGGTAATCCCAAAAGCGAC; RPL19-F: CCCATCTTTGATGAGCTTCC; RPL19- R: TGCTCAGGCTTCAGAAGAGG.

### Generation of precision-cut lung slices (PCLS) culture

For mouse PCLS, lung tissues were collected 8-10 weeks after LUAD induction in KPTI mouse. The lungs were perfused with PBS through the right ventricle and inflated with 1 to 2 ml of 2% agarose (Thermo Fisher, 16550100) dissolved in PBS by trachea. Lungs were dissected from the chest cavity and submerged in ice-cold PBS to solidify agarose. Lung lobes were sliced at a width of 500 µm using a vibratome (Leica, VT 1000S). The slices were cultured in DMEM/F-12 (Thermo Fisher, 11330032) with 1% Pen/Strep under standard cell culture conditions (37C, 5% CO2). ABT263 (2.5 µM), ABT737 (2 µM), Fisetin (10 µM), DQ (1 µM + 20 µM), and XL888 (1 µM) were treated during the culture. At day 5, cultured PCLSs were processed for downstream analyses.

### Flow cytometry analysis of mouse PCLS

The lung slices were placed into 15 ml conical tubes containing 1 ml of digestion cocktail of Dispase (3 U/ml, Thermo Fisher) and Dnase (50 U/ml, Sigma) after PBS washes. The slices were incubated in a digestion cocktail for 30 mins at 37°C with continuous shaking. The mixture was then washed with a FACS buffer (2% FBS and 1% Penicillin-Streptomycin in DMEM). The mixture was passed through a 70 μm cell strainer. Cells were stained with antibodies and analyzed by flow cytometry as described above.

### Micro-computed tomography (CT) data acquisition and analysis

An microCT system built for *in vivo* small animal imaging (U-CT, MILabs, Houten, The Netherlands) was used to measure in tumor volumes in vivo. During the scans, animals were maintained under anesthesia using approximately 2% isoflurane mixed with medical grade oxygen while a total of 1,440 projects were acquired over 360° with an x- ray tube voltage of 60 kVp and current of 0.24 mA. The projection data were acquired in a step-and-shoot mode with x-ray exposure time of 75 ms at each step, and there were two exposures at each step. No data binning was applied during the acquisition (i.e., 1×1 binning). Image reconstruction was performed using the vendor-provided conebeam filtered backprojection algorithm. The reconstructed image volumes were in the voxel size of 0.04 mm × 0.04 mm ×0.04 mm. The volumetric matrix sizes were dependent on the field of view selected during the reconstruction step focusing the lungs. The reconstructed CT images were imported to the software ITK-SNAP for lung tumor volume segmentation and measurement.

### Lentivirus infection

Primary human lung fibroblasts or mouse lung fibroblasts were seeded and infected the following day with Lenti-tTS/rtTA, Lenti-TRE-p16INK4a-T2A-dTomato, or Lenti-shApoe (CCTGAACCGCTTCTGGGATTACTCGAGTAATCCCAGAAGCGGTTCAGG). On day 1, the fibroblasts were infected with lentivirus at 5 multiplicity of infection (MOI) in DMEM-F12 with 10% FBS and polybrene at 5 µg/ml. On day 2, cells were washed with 4 times with PBS and then placed in regular media (DMEM-F12, 10%FBS, 1% PS).

Doxycycline (1 µg/ml) treatment began 72 to 96 hours later for Lenti-tTS/rtTA and Lenti-TRE-p16 dual-transduced cells.

### Fatty acid uptake analysis

We used BODIPY™ 500/510 C_1_, C_12_ (D3823) to assess the transfer of fatty acid into tumor organoids. After a two-week culture period, organoids were incubated overnight with 2 µg/ml of BODIPY-fatty acid. Following incubation, the Matrigel and organoids were dissociated to obtain single-cell suspensions. These single-cell suspensions were analyzed by flow cytometry, and BODIPY fluorescence intensity was measured to compare fatty acid transfer.

### Free fatty acid panel analysis

A pellet of approximately 200,000 cells were homogenized into 200uL of 10% methanol in water. A mix of deuterated fatty acids were spiked into 100uL of cell homogenate.

Samples were extracted using methanol and isooctane then derivatized using PFBB as previously described ^37^. Samples were analyzed by GC-MS on an Agilent 6890N gas chromatograph equipped with an Agilent 7683 autosampler. Fatty acids were separated using a 15m ZB-1 column (Phenomenex) and monitored using SIM identification.

Analysis was performed using MassHunter software.

### Survival analysis in TCGA LUAD

For survival analysis in TCGA LUADs, we uilized the web-based SurvivalGenie to generate the Kaplan-Meier plot^38^. The highly expressed genes from human CDKN2A+FAP+SMA+ cluster (Supplementary Table 9) were used as input. Patient tumors were categorized into high and low expression groups based on the median expression levels of these genes.

### Quantification and statistical analysis

GraphPad Prism was used for all statistical analyses. Statistical significance was determined by ordinary one-way ANOVA or a Student’s two-tailed unpaired t-test. For consistency in these comparisons, the following denotes significance in all figures: \**P* < 0.05, \*\**P* < 0.01, \*\*\**P* < 0.001, \*\*\*\**P* < 0.0001.

## Data availability

Previously published human scRNA-seq data that are re-analyzed in this study are available in NCBI Gene Expression Omnibus (GEO) under the accession number GSE183219. The sequencing data of the mouse that support the findings of this study have been deposited in the accession number GSE268478.

## Code availability

No custom codes were developed and used in this manuscript. All codes are available by request to the corresponding author.

