## Supplemental Information for "Senescent fibroblasts in the tumor stroma rewire lung cancer metabolism and plasticity"

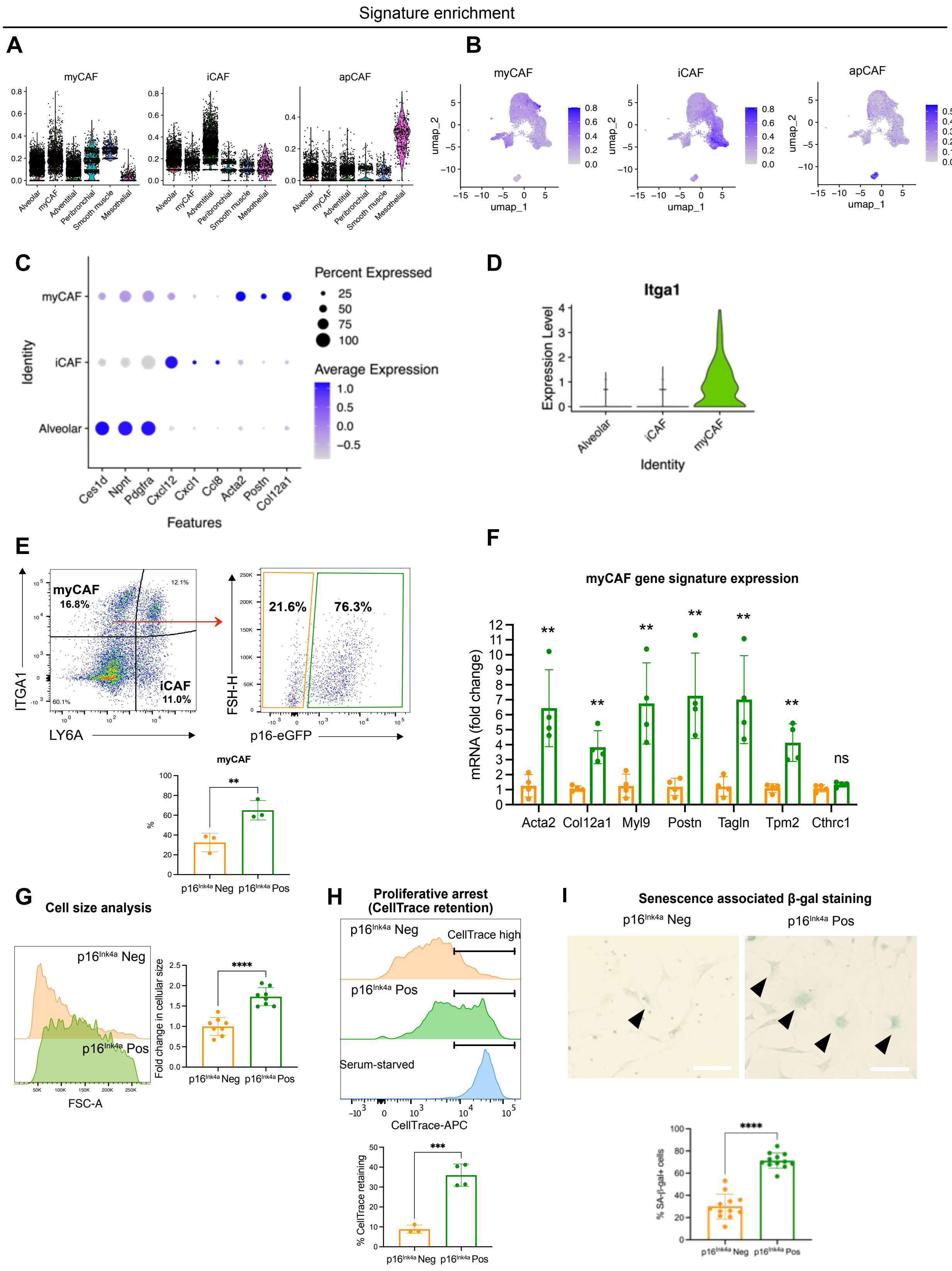

Signature enrichment
A
myCAF
iCAF
apCAF
Alveolar
myCAF
Adventitial
Peribronchial
Smooth muscle
Mesothelial
B
myCAF
iCAF
apCAF
umap_1
umap_2
C
Percent Expressed
25
50
75
100
Average Expression
1.0
0.5
0.0
−0.5
myCAF
iCAF
Alveolar
Identity
Ces1d
Npnt
Pdgfra
Cxcl12
Cxcl1
Ccl8
Acta2
Postn
Col12a1
Features
D
Itga1
Expression Level
Alveolar
iCAF
myCAF
Identity
E
myCAF
16.8%
12.1%
iCAF
11.0%
60.1%
ITGA1
LY6A
21.6%
76.3%
FSH-H
p16-eGFP
myCAF
**
%
p16Ink4a Neg
p16Ink4a Pos
F
myCAF gene signature expression
mRNA (fold change)
**
ns
Acta2
Col12a1
Myl9
Postn
Tagln
Tpm2
Cthrc1
G
Cell size analysis
p16Ink4a Neg
p16Ink4a Pos
FSC-A
Fold change in cellular size
****
H
Proliferative arrest
(CellTrace retention)
p16Ink4a Neg
CellTrace high
p16Ink4a Pos
Serum-starved
CellTrace-APC
% CellTrace retaining
***
I
Senescence associated β-gal staining
p16Ink4a Neg
p16Ink4a Pos
% SA-β-gal+ cells
****

**Fig S1. Senescent *p16$^{INK4a}$*+ fibroblasts contribute to myCAFs in mouse LUAD.**

(**A-B**) Gene signature enrichment was performed using gene lists from published myofibroblastic (myCAF), inflammatory (iCAF), and antigen-presenting (apCAF) cancer fibroblast datasets. (A) Vln plot of signature scores. (B) Visualization of signature scores on UMAP.

(**C**) Dot plots showing specific marker genes for each fibroblast cluster.

(**D**) Violn plot of Itga1 expression in myCAF cluster.

(**E**) Top: flow cytometry plot of GFP expression within ITGA1+ myCAF population. Bottom: Quantification of GFP+/GFP- fibroblasts within ITGA1+ myCAFs (n=3 mice for each group).

(**F**) qPCR analysis of myCAF genes in p16$^{Ink4a}$- and p16$^{Ink4a}$+ fibroblasts sorted from KPTI mouse lung (n=4 mice for each group).

(**G**) Quantification of p16$^{Ink4a}$- and p16$^{Ink4a}$+ fibroblast's cell size based on forward scatter on flow cytometry (n=8 mice for each group).

(**H**) Quantification of proliferative arrest based on CellTrace retention in p16$^{Ink4a}$- and p16$^{Ink4a}$+ fibroblast grown in proliferative conditions (n=3-4 mice for each group).

(**I**) Quantification of β-gal+ cell percentage in p16$^{Ink4a}$- and p16$^{Ink4a}$+ fibroblasts (n=12 for each group). Scale bars, 50 μm.

Unpaired *t*-test was used in (**E**), (**F**), (**G**), (**H**), (**I**) to test statistical significance. Data are represented as mean ± SD.; $^{**}P < 0.01$, $^{***}P < 0.001$, $^{****}P < 0.0001$

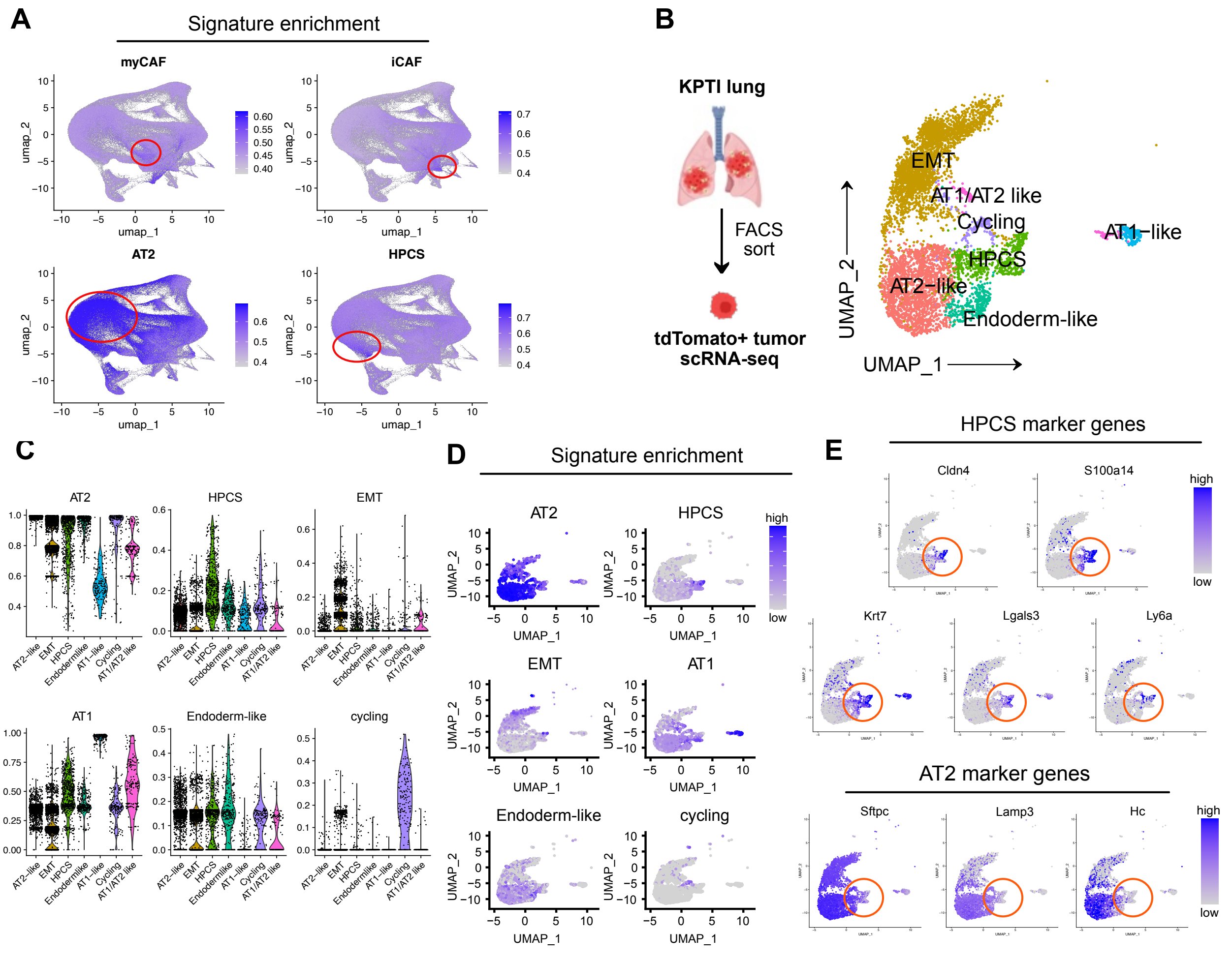


**Fig S2. Identification of HPCS cells in KPTI mouse lung.**

(**A**) Visualization of gene signature enrichment using previously published datasets of HPCS and CAFs in the UMAP clusters generated from spatial transcript identification on KPTI lung section.

(**B**) scRNAseq of tdTomato+ tumor cells isolated from KPTI mouse and unsupervised clustering of scRNA-seq data, annotated based on gene signature derived from Marjanovic et al. that described HPCS and other epithelial subtypes in LUAD.

(**C**) Violn plots of gene signature enrichment score of the epithelial subsets annotated in the scRNA-seq from KPTI lungs.

(**D**) Visualization of gene signature enrichment in UMAP.

(**E**) Feature plots showing HPCS marker and AT2 marker gene expressions.

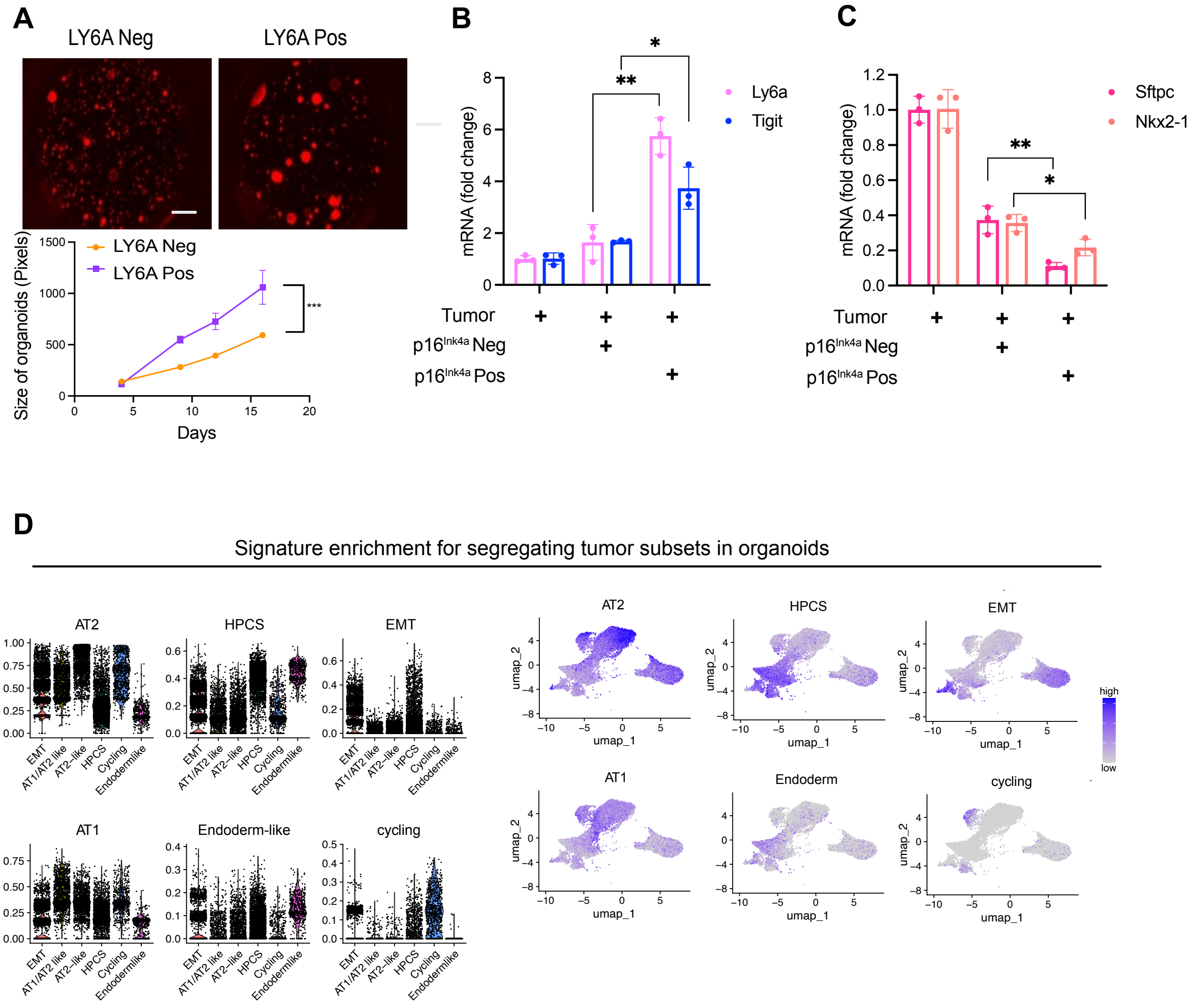


**Fig. S3.** ***p16**$^{Ink4a+}$* **CAFs promote LUAD progression *in vitro* by increasing HPCS cells.**

(**A**) Top: Tumor organoids using LY6A- and LY6A+ tumor cells isolated from KPTI mouse. Scale bars, 2000 μm. Bottom: Quantification of organoid size (n=4 for each group).

(**B**) qPCR analysis of HPCS genes in tdTomato+ tumor cells isolated from tumor organoids after co-culture (n=3 for each group).

(**C**) qPCR analysis of AT2 genes in tdTomato+ tumor cells isolated from tumor organoids after co-culture (n=3 for each group).

(**D**) Left: Violin plots of gene signature enrichment score in the scRNA-seq data from tumor organoids. Right: UMAP visualization of gene signature enrichment score in the scRNA-seq data from tumor organoids.

Unpaired *t*-test was used in (**A**), (**B**), and (**C**) to test statistical significance. Data are represented as mean ± SD.; $^*P < 0.05$, $^{**}P < 0.01$, $^{***}P < 0.001$

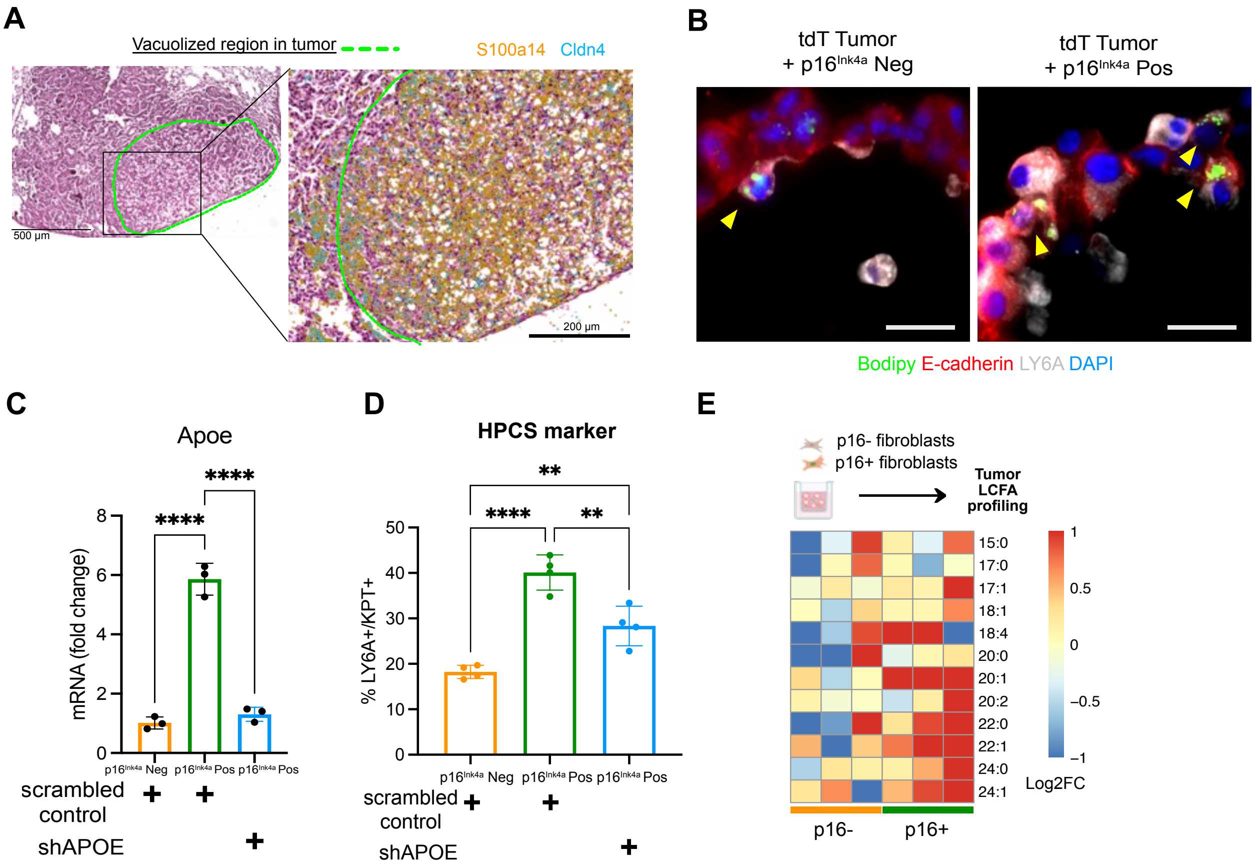


**Fig S4. APOE secreted from p16$^{Ink4a+}$ CAFs supports LUAD by regulating fatty acid transport.**

(**A**) Transcript mapping aligned with H&E staining demonstrates localization of HPCS genes within vacuolated regions in KPTI lung tissue.

(**B**) LY6A immunofluorescence with Bodipy 493/503 staining in tumor organoids after co-culture with *p16$^{Ink4a-}$* fibroblasts or *p16$^{Ink4a+}$* fibroblasts. Scale bars, 25 $\mu$m.

(**C**) Apoe transcript quantification in fibroblasts after down-regulation of Apoe using shRNA (n=3 for each group).

(**D**) Flow cytometry quantification of LY6A+ cells in tumor organoids co-cultured with *p16*$^{Ink4a-}$ fibroblasts or *p16*$^{Ink4a+}$ fibroblasts after down-regulation of Apoe (n=4 for each group).

(**E**) Heatmap depicting the log2 fold change of long chain free fatty acids profiled in tumor organoids co-cultured with either *p16*$^{Ink4a-}$ fibroblasts or *p16*$^{Ink4a+}$ fibroblasts.

One-way ANOVA was used in (**C**) and (**D**) to test statistical significance. Data are represented as mean ± SD.; $*P < 0.05$, $**P < 0.01$, $***P < 0.001$, $****P < 0.0001$

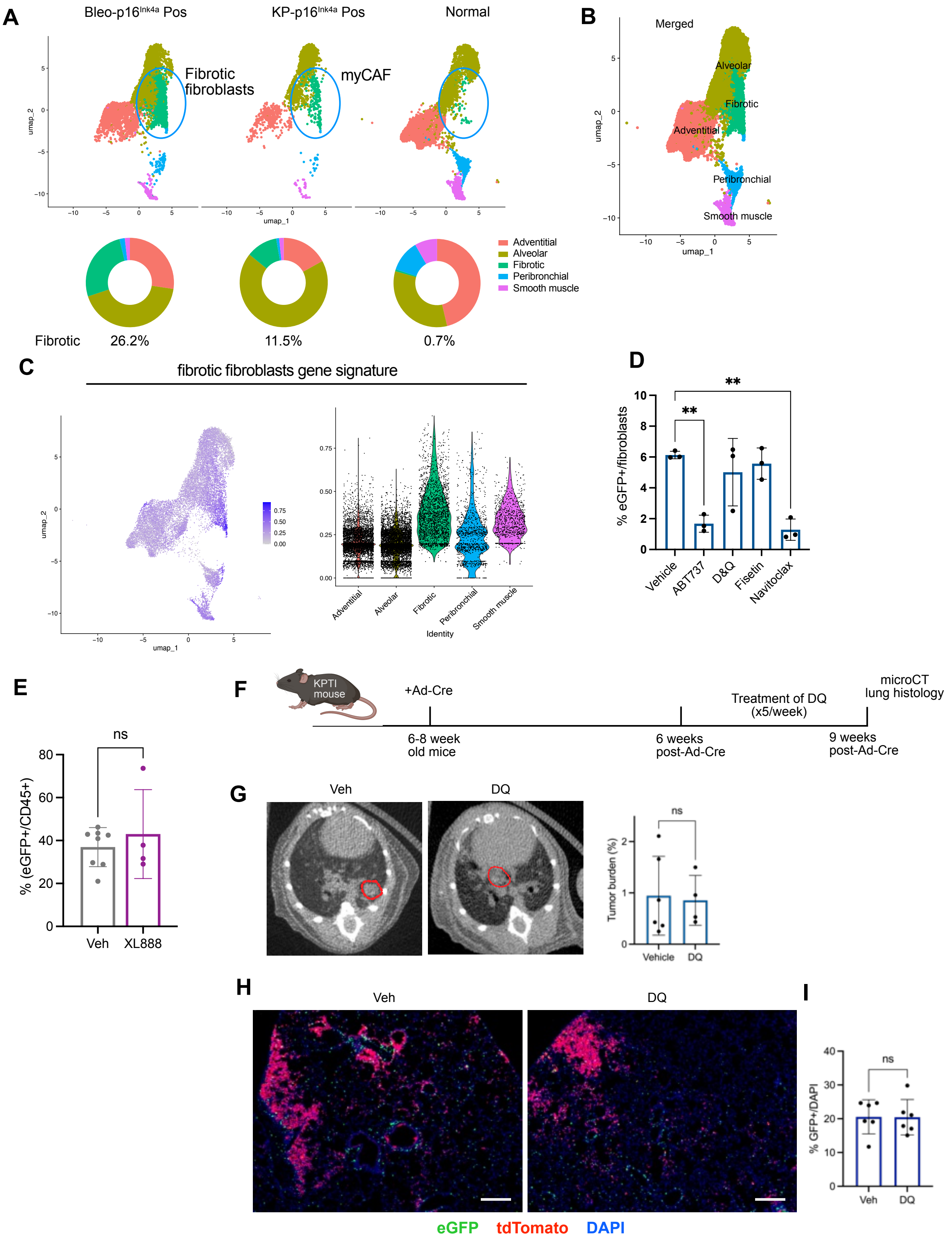


**Fig S5. Senolytic targeting of *p16^Ink4a+* CAFs**

(**A**) Top: UMAP plots of scRNA-seq data of *p16*$^{Ink4a+}$ fibroblasts isolated from bleomycin-injured lung, LUAD, and normal lung. Bottom: Proportion of fibroblasts within the fibrotic/myCAF fibroblast cluster relative to the total fibroblast population within each condition.

(**B**) Cell cluster annotation based on previously published single cell dataset of fibroblasts from fibrotic lungs (Tatsuya et al.).

(**C**) Fibrotic fibroblast gene signature enrichment analysis of scRNA-seq data established in (A).

(**D**) Percentage of GFP+ fibroblasts in PCLS after culture with indicated senolytics (n=3 slices for each group).

(**E**) Percentage of GFP+ cells within CD45+ immune cells in KPTI lung after XL888 administration (n=4-8 mice for each group).

(**F**) Experimental scheme of DQ treatment in KPTI mouse.

(**G**) Left: Representative image of microCT scan. Right: Quantification of lung tumor burden (n=4-6 mice for each group).

(**H**) Representative image of KPTI mouse lung after DQ treatment. Scale bars, 200 μm.

(**I**) Quantification of GFP+ cells in KPTI mouse lung sections (n=6 mice for each group).

Unpaired *t*-test was used in (**E**), (**G**), (**I**) and one-way ANOVA was used in (**D**) to test statistical significance. Data are represented as mean ± SD.; $^{**}P < 0.01$

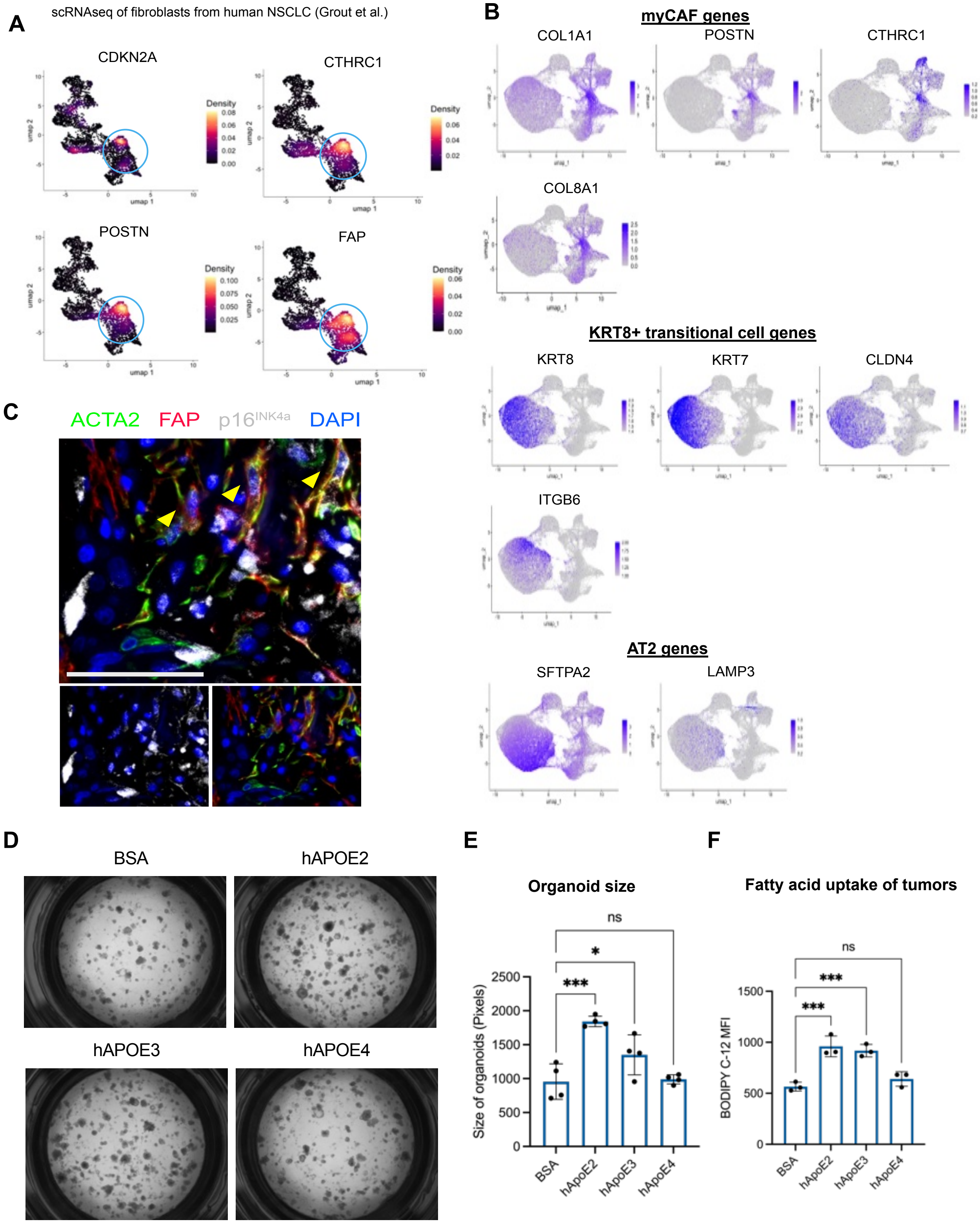


**Fig S6.** ***p16**$^{INK4A+}$* **CAFs and APOE in human LUAD.**

(**A**) Feature plots showing gene expressions of myCAF markers and CDKN2A in human NSCLC fibroblasts (analyzed from Grout et al.).

(**B**) Expression of myCAF, KRT8+ transitional cell, and AT2 genes in the UMAP clusters generated from spatial transcript analysis of human LUAD section.

(**C**) Representative immunofluorescence analysis of $p16^{INK4A}$ with ACTA2 and FAP in hLUAD. Scale bars, 50 $\mu$m.

(**D**) Representative images of hLUAD organoids cultured with normal human lung fibroblasts ad treated with human recombinant APOE variants. Scale bars, 2000 $\mu$m.

(**E**) Quantification of organoid size (n=4 for each group).

(**F**) Quantification of fatty acid transfer using Bodipy C-12 (n=3 for each group).

One-way ANOVA was used in (**E**) and (F) to test statistical significance. Data are represented as mean ± SD.; **P* < 0.05, ***P* < 0.01, ****P* < 0.001
